# Cross-Tissue Inflammatory Convergence Supports a Putative Muscle–Brain Axis in Alzheimer’s Disease

**DOI:** 10.64898/2026.10.07.751107

**Authors:** Andong Zhang, Wei Shen, Shijun Yan, Jing Chen, Xiaoyan Liu, Jingfeng Zou, Shaotian Li, Shanshan Hu, YiTing Liu, Catherine Pan, Zhuoqing Lan, Guoping Peng, Qin Zhang, Wen Peng, Fariba Ahmadizar, Jing Zhang, the Alzheimer’s Disease Neuroimaging Initiative

**Author notes:** Correspondence to: Jing Zhang, Department of Pathology, The First Affiliated Hospital, Zhejiang University School of Medicine, Hangzhou, Zhejiang, 311121, China., or Fariba Ahmadizar, Department of General Practice, Amsterdam University Medical Centre, Amsterdam, The Netherlands., or Wen Peng, Department of General Practice, Union Hospital, Tongji Medical College, Huazhong University of Science and Technology, No. 1277, Jie Fang Avenue, Wuhan, 430022, Hubei, China. Contributed equally (Andong Zhang, Wei Shen).

## Abstract

**Background:** Reduced muscle strength is associated with cognitive decline and Alzheimer’s disease (AD), suggesting that skeletal-muscle aging and AD-related neurodegeneration may share biological mechanisms. Vascular dysfunction, immune activation, and extracellular-matrix remodeling have been implicated in both tissues, but the extent of shared cellular and molecular programs remains unclear. We investigated cross-tissue transcriptional programs in aging skeletal muscle and AD-related brain tissue and evaluated whether related signals are detectable in peripheral blood.

**Methods:** We analyzed independent, unpaired human transcriptomic datasets from skeletal muscle (23 donors), the SEA-AD middle temporal gyrus atlas (84 donors), and peripheral blood. Donor-aware models identified age-associated changes in muscle and transcriptomic changes across continuous pseudoprogression score (CPS) in brain. Cross-tissue comparisons focused on 30 brain donors within an early CPS window. Shared tissue-derived programs were assessed in peripheral blood mononuclear cells and independently tested in frailty bulk-blood and Alzheimer’s Disease Neuroimaging Initiative (ADNI) whole-blood datasets. CellOracle simulations were used to prioritize candidate transcriptional regulators.

**Results:** Aging muscle showed increased inflammatory, extracellular-matrix, and stromal communication programs. During early AD-related brain progression, vascular and immune populations showed increasing inflammatory and matrix-associated activity together with shifts in intercellular communication. Cross-tissue analysis identified six concordantly enriched pathways comprising 109 leading-edge genes in endothelial populations and four pathways comprising 51 genes in myeloid populations. Related peripheral-blood signatures varied across cohorts and cell types. However, in independent frailty and ADNI mild cognitive impairment and AD datasets, 23 of 30 predefined tissue-derived pathway-subset tests showed significant positive enrichment after false-discovery-rate correction, whereas individual gene-level bulk-blood associations did not survive correction. CellOracle analyses prioritized ETS2 and BHLHE40 as exploratory endothelial regulators whose simulated perturbation consistently reduced selected shared program scores across tissues.

**Conclusions:** Aging skeletal muscle and AD-related brain tissue exhibit convergent endothelial and myeloid inflammatory programs, with related pathway-level signals detectable in peripheral blood. These cross-tissue programs and candidate endothelial regulators provide testable hypotheses for mechanisms linking systemic aging, muscle dysfunction, and AD-related neurodegeneration.

## BACKGROUND

Aging is accompanied by changes in skeletal muscle, the vasculature, the immune system and the brain[1]. Epidemiological studies link lower muscle strength with subsequent cognitive decline, including Alzheimer’s disease (AD), motivating investigation of a putative muscle–brain axis (MBA)[2,3]. At the tissue level, skeletal-muscle aging involves not only loss of muscle mass and function but also remodeling of stromal, vascular and inflammatory programs[4,5]. AD, in parallel, is increasingly understood as a multicellular disorder in which microglial activation, endothelial dysfunction, extracellular-matrix remodeling and altered vascular–immune interactions accompany neuronal pathology[6–8]. Whether aging skeletal muscle and AD-related brain tissue share molecular programs at cell-type resolution, however, remains unclear.

Existing muscle-aging and AD atlases allow comparison of transcriptional programs across tissues and assessment of related signals in circulating immune cells. A key question is whether shared pathway labels reflect overlapping genes within corresponding endothelial and myeloid populations. Inflammatory, cytokine-response and extracellular-matrix pathways recur across many age-related conditions, so shared labels alone are insufficient to establish a specific muscle–brain mechanism, common cellular origin or direction of signaling[1].

Endothelial cells provide a biologically relevant compartment for this comparison because they are represented in both skeletal muscle and brain and participate in inflammatory signaling, leukocyte interactions and tissue homeostasis[7,9]. Stromal, myeloid and other immune populations may contribute related programs within each tissue, placing endothelial responses within a broader multicellular context. Peripheral blood offers a complementary systemic readout of these programs, although their detection in circulating immune cells does not itself establish mediation between muscle and brain[10].

Here, we compared independent, unpaired human skeletal-muscle, brain and peripheral-blood transcriptomic datasets to investigate a putative MBA in AD. We first characterized age-associated transcriptional and inferred communication changes in muscle, alongside brain changes associated with continuous pseudoprogression score (CPS). We then identified shared endothelial and myeloid programs, examined related patterns in peripheral blood mononuclear cells (PBMCs), and tested tissue-derived pathway subsets in an independent frailty bulk-blood cohort and in mild cognitive impairment (MCI) and AD comparisons from the Alzheimer’s Disease Neuroimaging Initiative (ADNI). Finally, we used CellOracle simulations of transcription-factor (TF) knockout in endothelial and myeloid populations to prioritize candidate regulators of the shared programs[11].

## METHODS

### Overall study design and levels of evidence

We performed secondary analyses of independent public skeletal-muscle, brain and peripheral-blood transcriptomes. HLMA age-group comparisons and SEA-AD CPS associations defined tissue-specific references (Figures 1–2). We then examined shared inflammatory pathways and leading-edge genes in endothelial and myeloid populations (Figures 3–4), followed by assessment of related immune programs and tissue-derived genes in independent PBMC cohorts (Figure 5) and external evaluation of the tissue-intersection genes and pathway subsets in the GSE206762 frailty and ADNI MCI/AD bulk-blood comparisons (Supplementary Figures S6–S8). CellOracle simulated cluster-specific knockout of tissue-derived endothelial and myeloid TF candidates to evaluate predicted cross-tissue program and gene responses (Figure 6; Supplementary Figures S9–S10). Tissues were not sampled from matched individuals, and cross-tissue convergence denoted concordant program directions rather than transmission, mediation or disease-level causality.

**Fig. 1.**
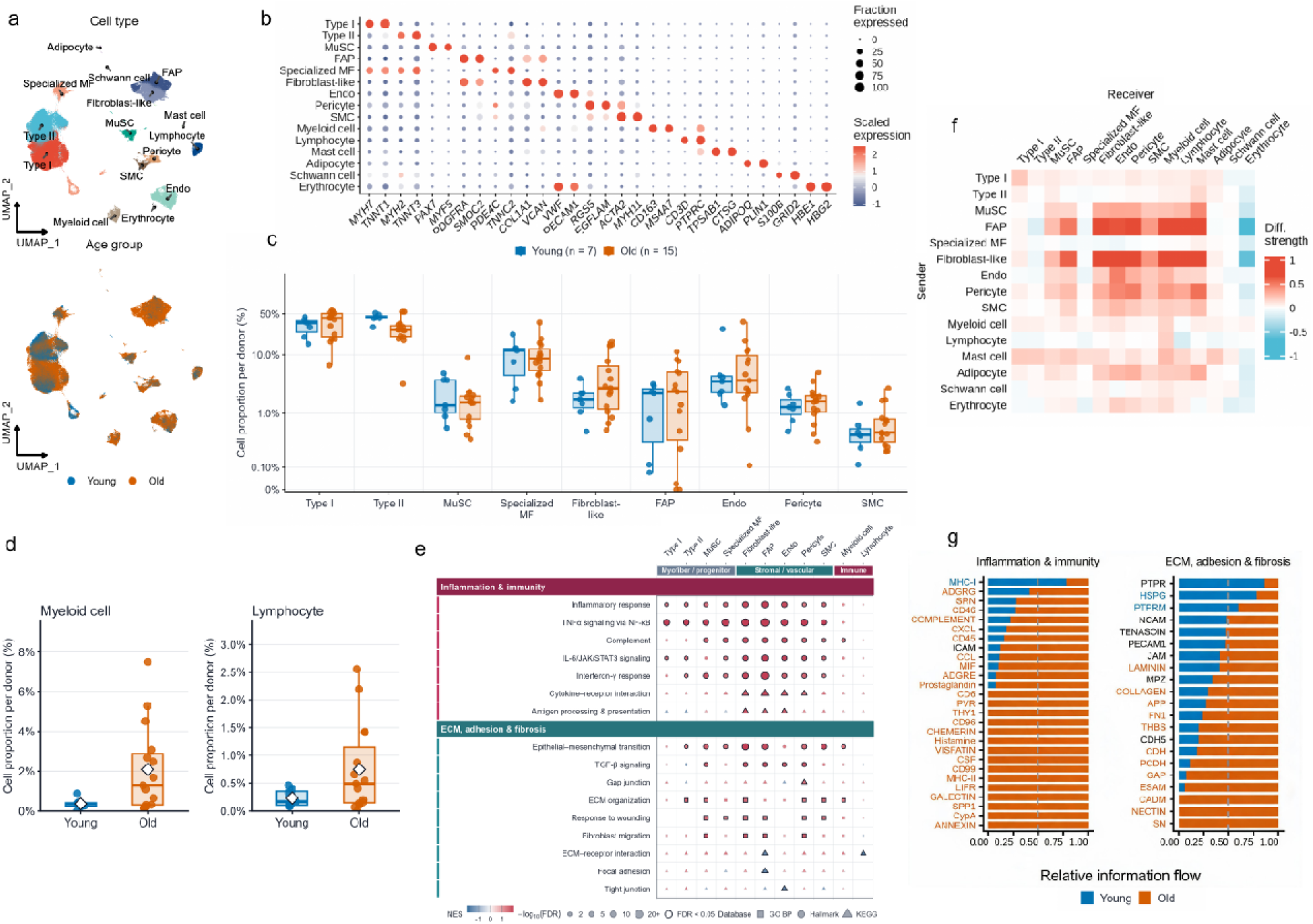
Cellular composition and inflammatory programs in aging skeletal muscle. (a, b) Cell-type and age-group UMAPs and a 30-gene marker dot plot for the human limb skeletal muscle atlas, comprising 292,423 cells or nuclei from 23 donors (8 Young and 15 Old). Dot size denotes the expressing fraction and color the gene-scaled mean expression. (c, d) Donor-level proportions in the common-technology single-nucleus subset (7 Young and 15 Old donors). The denominator was all annotated nuclei per donor. Points represent donors; boxes show medians and interquartile ranges; whiskers extend to 1.5 times the interquartile range; white diamonds in d denote means. Panel c uses a pseudo-log percentage axis and d separate linear axes. Two-sided Wilcoxon rank-sum tests were adjusted across 15 populations by the Benjamini–Hochberg method. (e) Gene set enrichment analysis (GSEA) of 7 inflammatory or immune and 9 extracellular-matrix, adhesion, or fibrosis pathways across 11 cell types. Rankings were derived from donor-aggregate limma-trend models. Positive normalized enrichment scores (NESs) favor Old; dark outlines denote original full-family false discovery rate (FDR) < 0.05. Point size denotes −log10(FDR); shapes distinguish Hallmark, KEGG and GO BP collections. Blank cells were unavailable. (f) Old-minus-Young differences in CellChat weighted communication strength, with senders in rows and receivers in columns. (g) Group-pooled relative information flow for 48 pathways; Young and Old shares sum to 1 within each pathway. Panels f and g are model-based pooled summaries rather than independent-donor tests. FAP, fibro/adipogenic progenitor; MuSC, muscle stem or satellite cell; ECM, extracellular matrix.

**Fig. 2.**
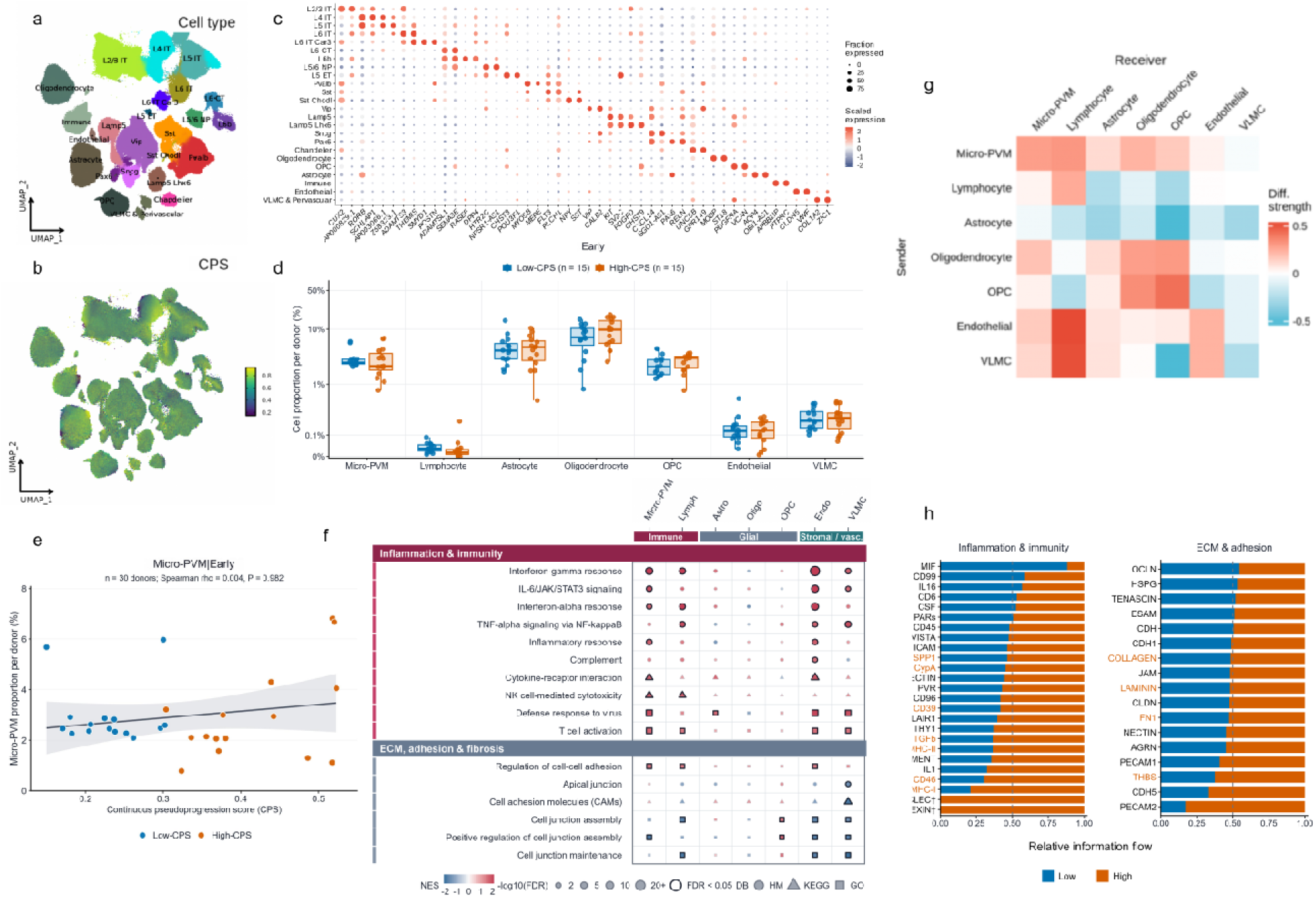
Brain programs and inferred communication in the early CPS window. (a–c) Cell-type and continuous pseudoprogression score (CPS) UMAPs and a 48-gene marker dot plot for the SEA-AD middle temporal gyrus atlas (84 donors; 1,178,694 nuclei). Dot size denotes the expressing fraction and color the gene-scaled mean expression. (d) Donor-level proportions of seven non-neuronal populations in the early window (CPS ≤ 0.55), after exclusion of source-defined severely affected donors. The CPS median (0.3030) defined Low- and High-CPS groups (15 donors each). Points represent donors; boxes show medians and interquartile ranges; the percentage axis uses a pseudo-logarithmic scale. Two-sided Wilcoxon rank-sum tests used continuity correction and Benjamini–Hochberg adjustment across seven populations. (e) Microglia/perivascular macrophage (Micro-PVM) proportion versus continuous CPS in 30 donors. The reported Spearman correlation is two-sided; the line and 95% confidence band are an unadjusted descriptive linear fit. (f) GSEA of 10 inflammatory or immune and six extracellular-matrix, adhesion, or fibrosis pathways across seven populations. NESs were obtained from continuous-CPS rankings generated by donor-random-effect NEBULA models; positive values indicate enrichment with increasing CPS. Dark outlines denote original full-family FDR < 0.05. Models included 30 donors, except Lymphocyte (28 donors). The VLMC enrichment unit pooled the VLMC & Perivascular subclass. (g) CellChat High-minus-Low differences in weighted communication strength. (h) Relative information flow for 42 pathways; shares sum to 1 within each pathway. Among pathways detected in both groups, 11 passed the ligand–receptor-record-level FDR threshold and all had greater relative flow in the High-CPS group. ANNEXIN and CLEC were detected only in the High-CPS network and were summarized descriptively rather than included in the testing family. Two-sided paired Wilcoxon tests compared matched ligand–receptor records, with adjustment across 96 pathways with valid tests. Panels g and h are pooled network inferences rather than donor-level tests. OPC, oligodendrocyte precursor cell; VLMC, vascular and leptomeningeal cell.

**Fig. 3.**
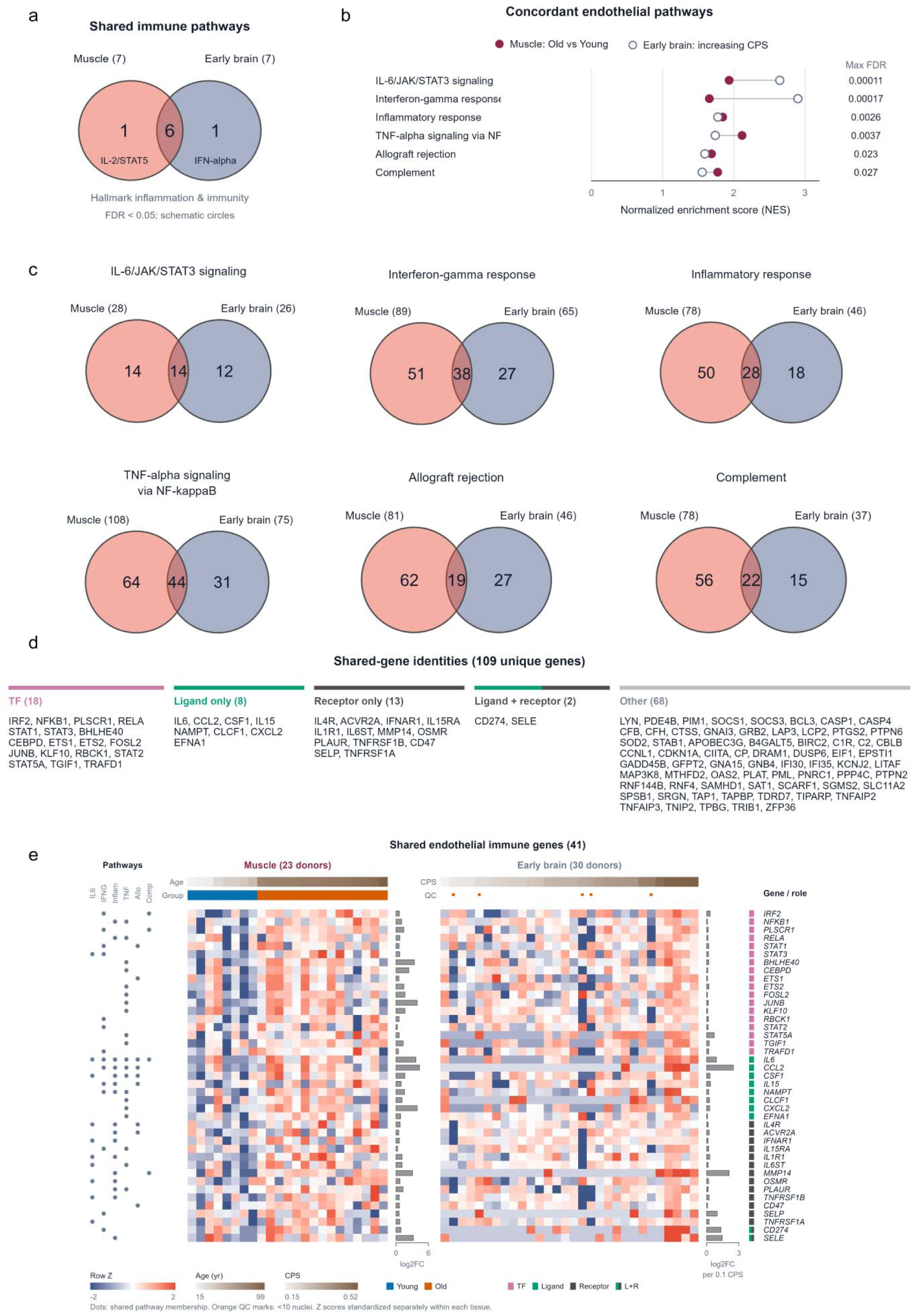
Shared endothelial inflammatory programs across muscle and brain. (a) Intersection of significant pathways within a prespecified eight-pathway Hallmark inflammatory or immune module. Seven pathways met the original FDR < 0.05 threshold in each tissue, with six shared. (b) NESs for the six concordant pathways. Muscle estimates compare Old with Young; brain estimates represent continuous CPS associations. Lines pair tissue-specific estimates for the same pathway. The displayed maximum FDR is a ranking statistic, not a combined probability. (c) Venn diagrams of complete native GSEA leading-edge sets. Muscle-only, shared, and brain-only counts are shown; shared counts were 14, 38, 28, 44, 19, and 22 in the order displayed. Circle area is not proportional to set size, and overlap significance was not tested. (d) Roles of the 109 genes shared across tissues and deduplicated across pathways: 18 transcription factors (TFs), 8 ligand-only genes, 13 receptor-only genes, 2 ligand–receptor genes, and 68 other genes. Roles were assigned from HumanTFs and CellChatDB and do not demonstrate functional activity. (e) Donor heatmaps for the 41 TF, ligand, or receptor genes. Muscle included 23 donors (8 Young, 15 Old; 23,227 endothelial cells), and brain included 30 donors (693 endothelial nuclei). Gene-wise Z scores were calculated separately within each tissue; columns were ordered by age or CPS. Five brain donors with fewer than 10 endothelial nuclei are flagged. Side bars show muscle limma log2 fold changes and brain NEBULA log2 fold changes per 0.1 CPS and are not directly comparable across tissues. The complete 109-gene set is shown in Supplementary Fig. S4.

**Fig. 4.**
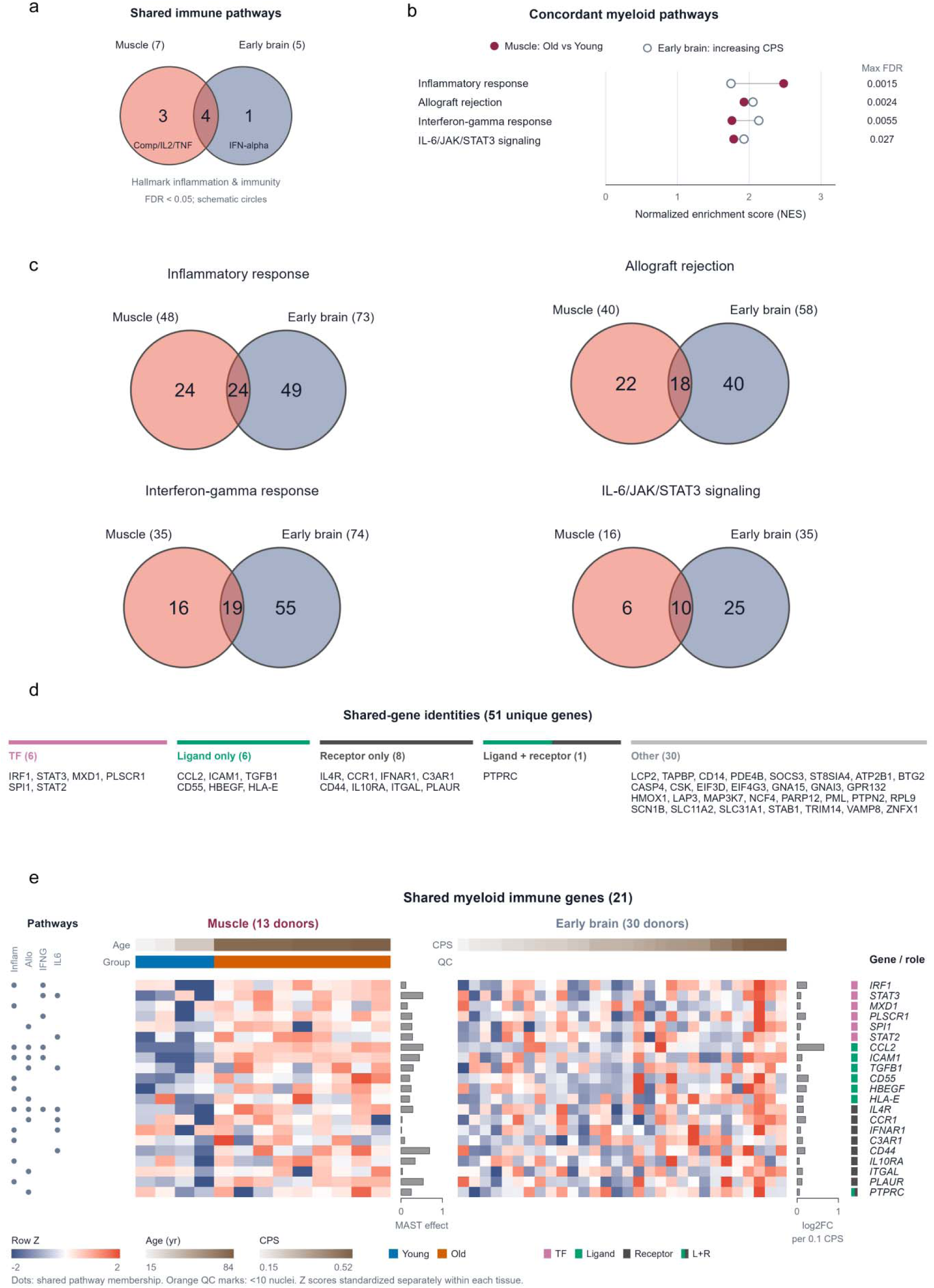
Shared myeloid inflammatory programs across muscle and brain. (a) Intersection of significant pathways within the prespecified eight-pathway Hallmark inflammatory or immune module. Seven muscle and five brain pathways met the original FDR < 0.05 threshold, with four shared. (b) NESs for the four concordant pathways. Muscle estimates compare Old with Young; brain Micro-PVM estimates represent continuous CPS associations. Lines pair estimates for the same pathway. The displayed maximum FDR is a ranking statistic rather than a combined probability. (c) Complete leading-edge intersections for inflammatory response, allograft rejection, interferon-γ response, and IL-6/JAK/STAT3 signaling. Total muscle, total brain, and shared leading-edge gene counts were 48/73/24, 40/58/18, 35/74/19, and 16/35/10, respectively. Circle area is not proportional to set size, and overlap significance was not tested. (d) Roles of the 51 unique shared genes: 6 TFs, 6 ligand-only genes, 8 receptor-only genes, 1 ligand–receptor gene, and 30 other genes. Annotations do not establish protein secretion, TF activity, or receptor activation. (e) Donor heatmaps for 21 annotated genes. Muscle included 4 Young donors (259 nuclei) and 9 Old donors (4,166 nuclei); brain included 30 donors (13,416 Micro-PVM nuclei). Within-tissue gene Z scores are shown. Muscle side bars represent MAST-adjusted predicted mean log2-expression differences; brain bars represent NEBULA log2 fold changes per 0.1 CPS. The complete 51-gene set is shown in Supplementary Fig. S5. Donors are the biological replicates.

**Fig. 5.**
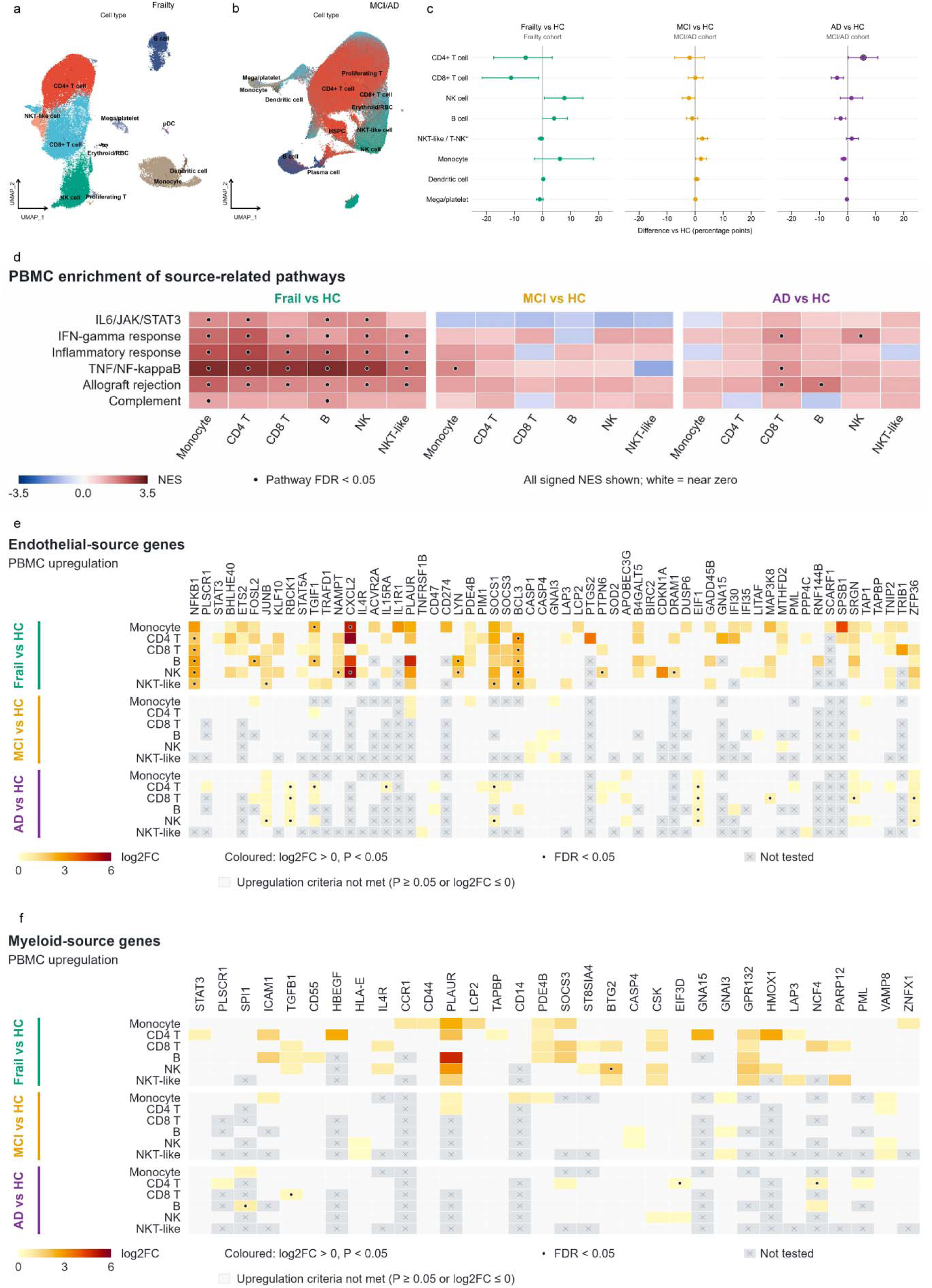
Peripheral immune representation of tissue-derived inflammatory programs. (a) GSE157007 UMAP of 76,509 peripheral blood mononuclear cells (PBMCs) from six healthy older controls and five participants with source-defined Frailty. (b) SRP312418 atlas of 696,093 PBMCs from healthy controls (HC), mild cognitive impairment (MCI), and Alzheimer disease (AD); the UMAP shows a diagnosis- and cell-type-stratified subset of 250,096 cells. Frailty denotes the source phenotype rather than diagnosed sarcopenia; the atlases were not jointly embedded. (c) Participant-level mean differences in cell proportions relative to cohort-specific controls. Points show mean percentage-point differences and bars bootstrap 95% confidence intervals. Analyses included 121 HC, 46 MCI, and 27 AD participants and, independently, six HC and five Frailty participants. Black outlines denote effects retained by scCODA at 5% estimated false discovery rate (FDR); only the AD-associated increase in CD4+ T cells met this criterion. (d) Gene set enrichment analysis of six source-related Hallmark pathways across monocytes, CD4+ T, CD8+ T, B, NK, and NKT-like cells in Frailty versus HC, MCI versus HC, and AD versus HC. Color denotes normalized enrichment score; positive values favor the case group. Dots denote FDR < 0.05 after Benjamini-Hochberg correction across eligible pathway-cell-type tests within each contrast. (e, f) Projection of the complete 109-gene endothelial and 51-gene myeloid tissue intersections into PBMC differential-expression results. Genes were retained when log2 fold change was positive and unadjusted P < 0.05 in at least one displayed context, yielding 61 endothelial-source genes in e and 32 myeloid-source genes in f. Colored tiles denote comparison-specific positive nominal results; pale entries denote tested results that did not meet this display rule, crosses unavailable tests, and dots gene-level FDR < 0.05 from the complete testing family. Source labels indicate provenance and do not assign a circulating cell of origin.

**Fig. 6.**
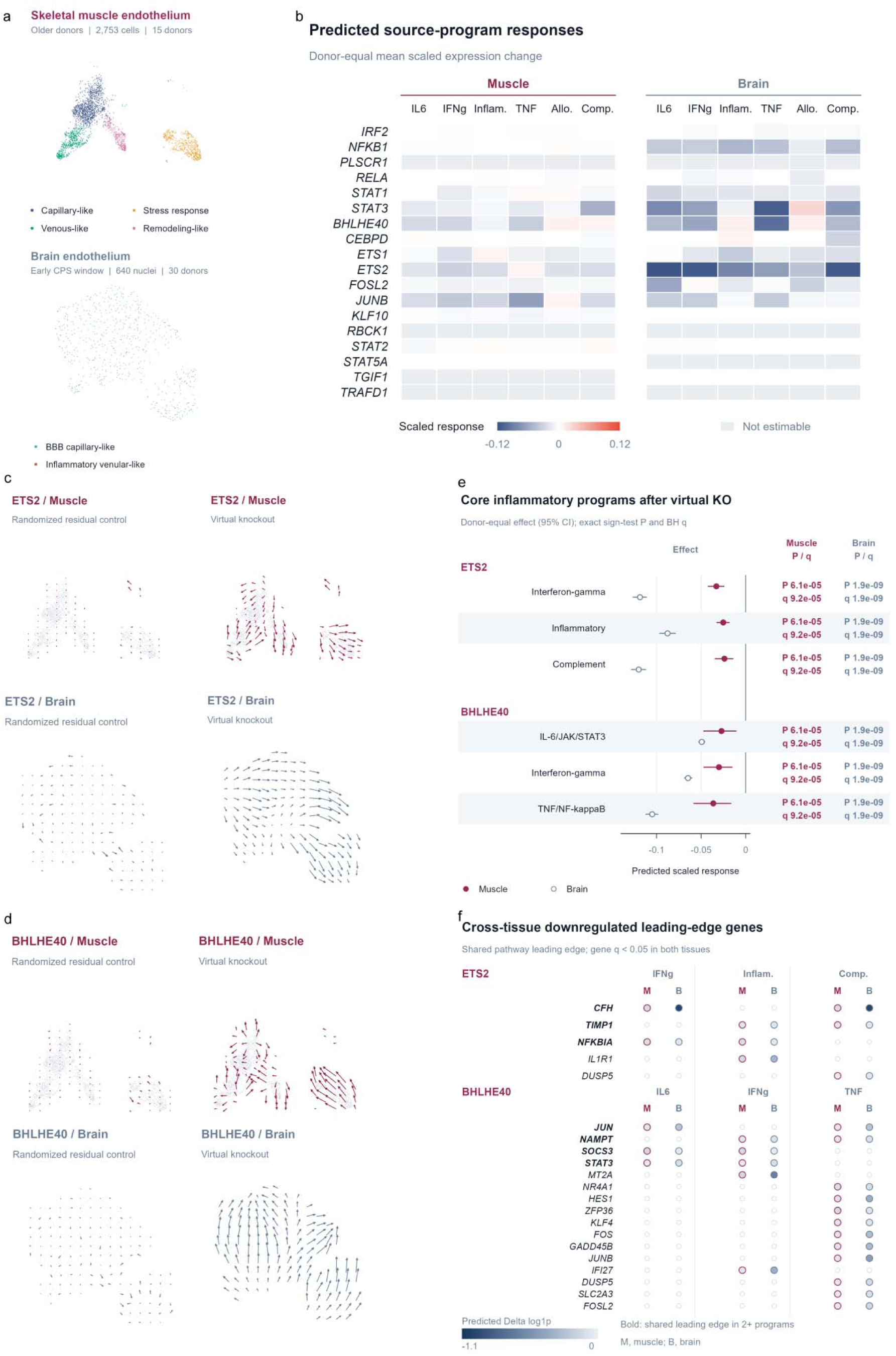
Endothelial ETS2 and BHLHE40 virtual-knockout responses across muscle and brain. (a) Endothelial state spaces used for perturbation modeling. Skeletal muscle comprised 2,753 cells from 15 older donors and four marker-informed descriptive states; brain comprised 640 nuclei from 30 early-window donors and two descriptive states. State labels aid interpretation and do not define a new taxonomy or trajectory. (b) Donor-equal responses of six source inflammatory programs after virtual knockout (KO) of 18 TF candidates derived from the Fig. 3 intersection. Color denotes the standardized KO response; gray denotes a non-estimable combination. (c, d) CellOracle transition-vector fields for ETS2 and BHLHE40 KO, respectively, in muscle and brain, each with a randomized-residual control. Arrows show local model-predicted changes rather than measured cell movement or exact KO endpoints. Scaling is shared between KO and control within tissue, but not across tissue-specific state spaces. (e) Predicted effects on selected core programs: interferon-γ response, inflammatory response, and complement for ETS2; and IL-6/JAK/STAT3, interferon-γ response, and TNF-α/NF-κB signaling for BHLHE40. Points show donor-equal means and bars donor-bootstrap 95% intervals. Two-sided exact sign tests used donor response directions; Benjamini–Hochberg q values were calculated across six source programs within each KO and tissue. All 15 muscle and 30 brain donors showed negative predicted directions. (f) Cross-tissue genes that were leading-edge members in both tissues, had negative predicted gene effects, and had gene-level q < 0.05 in both tissues. Bold genes occurred in at least two selected programs. Source-program GSEA remained nonsignificant after global correction (all q > 0.97); these are directional exploratory candidates. CellOracle perturbations are model predictions and do not establish causal rescue.

Additional file 1 contains the Supplementary Methods and Supplementary Table S1. Additional file 2 contains Supplementary Figures S1–S10 and their legends. Additional file 3 is an Excel workbook containing pathway-selection records, analysis diagnostics and supporting source tables (Supplementary Tables S2–S6).

### Human skeletal muscle atlas analysis (Figure 1)

#### Study design and data source

This study was a secondary analysis of the publicly released human limb skeletal muscle single-cell and single-nucleus atlas[4]. The primary exposure was the source-defined age group, comparing 15 Old donors aged 77–99 years with 8 Young donors aged 15–45 years. All 23 donors represented in the analyzed released object were included, with no prospective sample-size calculation. Median ages were 82.0 and 31.5 years, respectively (Supplementary Table S1). The cached Seurat input contained 292,423 cells and nuclei and 48,355 RNA features; the myeloid MAST analysis used a subset of this same object. The AnnData X matrix contained log-normalized expression values; raw counts were not available. We converted the h5ad object to a Seurat v5 object with h5ad_to_srt() from the scop package and retained the atlas annotations for 15 cell types[12]. Sex and country were included as donor-level covariates; both sexes and both countries were represented in each age group (older: 8 female and 7 male donors, 9 from China and 6 from Spain; young: 3 female and 5 male donors, 5 from China and 3 from Spain). Cells labeled “Tenocyte” in the released object were renamed fibroblast-like cells, consistent with the source study and their marker profile (*THY1*, *CXCL14*, *PRG4*, *COMP*, and *FMOD* positive; *SCX*, *TNMD*, and *MKX* negative).

#### Cell-type annotation assessment

Data-driven markers were identified for each cell type using the cell-level Wilcoxon test implemented in RunDEtest (scop). This analysis was used only to assess the supplied annotations and was not interpreted as donor-level differential expression. Canonical markers were retained when they ranked among the top 15 data-driven markers and the difference in the fraction of expressing cells between the target population and all other cells exceeded 0.1. Otherwise, replacement markers were selected by an effect-size score defined as log2 fold change multiplied by the difference in expressing-cell fraction. No more than two markers were retained per cell type. Ribosomal protein genes, mitochondrial genes, pseudogenes, and unnamed transcripts were excluded.

#### Donor-level cell-composition analysis

Cell-composition analysis was restricted to snRNA observations to use a common capture technology across age groups (212,774 nuclei; Young, n = 7 donors; Old, n = 15 donors). Donor, rather than nucleus, was the biological replicate. For each donor, the proportion of each of the 15 annotated cell types was calculated relative to all annotated snRNA nuclei from that donor; donor-by-cell-type combinations with no observed nuclei were retained as zero. Young and Old donor proportions were compared separately for each cell type using a two-sided Wilcoxon rank-sum test without continuity correction, and P values were adjusted across the 15 tests with the Benjamini-Hochberg method[13]. Figure 1c displays raw donor percentages as points with Tukey-style box plots and uses a base-10 pseudo-logarithmic y-axis transformation (sigma = 0.05 percentage points) that retains zero values. Figure 1d focuses on myeloid cells and lymphocytes, using untransformed linear percentage scales with separate y-axis ranges; white diamonds indicate arithmetic means. All tests were performed on the original percentages. Sensitivity analyses included technology-aggregated proportions for all 23 donors and an equal-donor-weight linear model of arcsine-square-root-transformed proportions adjusted for country and sex.

#### Donor-level differential expression using aggregated normalized expression

Donors were the biological replicates for differential-expression inference[14,15]. Because raw counts were unavailable, log-normalized values were back-transformed with expm1 and summed across cells within each donor-by-cell-type unit to generate aggregated normalized-expression values. This procedure did not recover the original molecule counts. Units containing fewer than 20 cells were excluded. Genes were retained if their summed expression reached at least 10 in at least 3 retained units, and each age-group comparison required at least 3 donors per group.

For each retained dataset, an edgeR DGEList was created, trimmed mean of M-values (TMM) normalization factors were estimated, and normalized values were transformed to log2 counts per million (CPM) with a prior count of 1[16,17]. Differential expression was modeled with limma-trend using ∼ age_pop + Sex + Country, followed by eBayes(trend = TRUE, robust = TRUE)[18]. The contrast was older versus young; positive coefficients therefore indicated higher expression in older donors. Benjamini-Hochberg correction was applied across tested genes separately within each dataset, with a false discovery rate (FDR) below 0.05 as the primary threshold. Nominal *P* values were additionally evaluated by 1,000 country-stratified permutations as supporting calibration. Age-group labels were permuted between donors within country, holding expression, sex and country fixed; the model ∼ age_pop + Sex + Country was refitted with robust limma-trend moderation for each permutation. Two-sided empirical P values were (1 + the number of permuted absolute moderated t statistics at least as large as observed)/1,001; permutation b used R seed b. The primary analysis covered eight major cell types and an overall aggregate (nine datasets). Pericytes, SMCs, and lymphocytes had fewer evaluable donor-level units and were analyzed separately as lower-powered, directional evidence.

#### Gene set enrichment analysis

Within each cell type, genes were ranked in descending order by the moderated t statistic from the donor-level aggregated-expression model without imposing a differentially expressed gene threshold. For duplicated gene symbols, the maximum signed t statistic was retained. Gene set enrichment analysis (GSEA) was performed with clusterProfiler::GSEA using the fgseaMultilevel engine (minimum gene-set size = 10; maximum gene-set size = 500; eps = 1e-50; seed = 42)[19,20]. The displayed analyses used four local Molecular Signatures Database (MSigDB) v2026.1 collections: Hallmark, KEGG_legacy, Gene Ontology Biological Process (GO_BP), and GTRD transcription factor targets (TFT_GTRD)[21–24]. Benjamini-Hochberg correction was applied separately across eligible gene sets within each dataset and collection. GO_BP terms were reduced for redundancy using Wang semantic similarity at a threshold of 0.7[25]. GSEA was repeated with log2 fold-change-ranked genes as a sensitivity analysis. The curated display pool comprised 33 pathways that reached FDR < 0.05 in at least one cell type after cross-collection semantic deduplication while retaining representative positive and negative signals. Figure 1e displays a focused subset of 16 pathways: seven inflammation/immunity pathways and nine ECM/adhesion/fibrosis pathways; the remaining 17 signaling/stress-response and metabolism pathways were retained in the source results but are not displayed in the manuscript. Unavailable pathway-by-cell-type combinations were left blank without imputation. The 33-pathway curated pool, biological themes and final display membership are provided in Supplementary Table S2a (Additional file 3).

#### Cell-cell communication analysis

Separate CellChat v2 networks were estimated from cells pooled within Young (86,588 cells) and Old (205,835 cells) groups using CellChatDB.human and the trimean estimator[26,27]. Old-minus-Young differences in weighted sender–receiver communication strength are shown in Fig. 1f. Interaction counts followed CellChat’s internal inference criterion; implementation settings are provided in Supplementary Methods.

Pathway-level comparisons used CellChat rankNet. Relative information flow denotes normalized between-group shares within each pathway; it does not compare absolute flow between pathways. CellChat compared sender–receiver cell-type-pair weights using paired Wilcoxon tests. For pathways detected in only one group, the software-assigned P = 0 was a detection convention, not a valid inferential P value. These pathways were interpreted descriptively; source label colors were not treated as evidence of BH-adjusted or donor-level significance. The union contained 84 pathways. TNF was excluded from presentation because its ligand transcript was below the detection limit in the older group. Pathways were functionally curated using CellChatDB structural annotations, manual review, and Gene Ontology enrichment; for the main-figure display, Fig. 1g was restricted to 48 pathways: 27 inflammation/immunity pathways and 21 ECM/adhesion/fibrosis pathways. Other annotated categories were not displayed or interpreted in the main figure. Robustness was assessed by repeating the complete workflow after downsampling to 1,000 cells per cell type per group with a fixed random seed.

#### Statistical interpretation and reporting

Donors were the biological replicates for composition and differential-expression models; GSEA used gene rankings from the corresponding donor-level models. Cell-level Wilcoxon tests were restricted to annotation assessment. CellChat pooled cells within age groups and therefore provided descriptive model-based comparisons rather than donor-level hypothesis tests. Limited donor representation reduced power for pericytes, SMCs and lymphocytes; nonsignificant findings were not interpreted as no change. The absence of raw counts constrained inference to aggregated, back-transformed normalized expression analyzed with limma-trend. The primary composition analysis used the shared snRNA-seq technology, whereas source expression and communication analyses included single-cell and single-nucleus data. Restricting composition analysis by technology did not constitute an snRNA-seq-only validation of the expression models. Figure 1 describes age-associated differences in the source atlas, without assuming that Old donors had clinically diagnosed sarcopenia.

### SEA-AD middle temporal gyrus analysis (Figure 2)

#### Data source, cohort, and CPS windows

We analyzed single-nucleus RNA-sequencing data from the SEA-AD MTG atlas, using the final-nuclei object released on June 22, 2026. The full cohort comprised 84 donors aged 65–102 years (51 female and 33 male; median age, 90 years) and 1,178,694 nuclei, used for atlas visualization and marker displays. Donor covariates and CPS were joined by donor identifier, and official UMAP coordinates were retained. Inferential analyses were restricted to 73 donors not designated as severely affected in the source metadata (43 female and 30 male; aged 69–102 years; median age, 91 years). The source study described the excluded 11 donors as a distinct, severely affected group with reduced nuclear RNA and chromatin accessibility, potentially reflecting advanced pathology rather than solely technical sample failure[28]. This restriction limits inference to donors outside that source-designated state. All 11 excluded donors had high AD neuropathological change and CPS values of 0.729–0.924, outside the early window. Donor characteristics are summarized in Supplementary Table S1.

The early CPS window was defined as CPS ≤ 0.55 (30 donors aged 72–102 years), and the late window as CPS > 0.45 (48 donors aged 69–100 years). Five transition-range donors contributed to both local analyses; the windows were not independent groups. The early window contained 25 donors recorded as having no dementia and five with dementia. These analytical cutoffs delineate local ranges of neuropathology-derived cross-sectional ordering, not validated clinical stages or an equivalent of MCI[28]. Excluding severely affected donors did not alter early-window membership, but this does not establish robustness to alternative CPS boundaries.

#### Cell annotation and analysis units

Official SEA-AD class, subclass, and supertype annotations were retained. To document how the immune compartment mapped onto the downstream analysis units, immune nuclei were displayed at supertype resolution. Monocyte comprised only 12 nuclei and was excluded from the immune UMAP, marker display, and pooled Micro-PVM unit. The retained immune compartment contained 36,674 nuclei assigned to six Micro-PVM supertypes and Lymphocyte. Twelve canonical identity markers were cross-checked against data-derived supertype marker rankings and visualized across the seven retained immune supertypes (Supplementary Figure S2a,b). The seven downstream analysis units were Micro-PVM, Lymphocyte, Astrocyte, Oligodendrocyte, OPC, Endothelial, and VLMC. Micro-PVM pooled the six retained Micro-PVM supertypes, whereas Lymphocyte was analyzed at the supertype level; the remaining units were analyzed at the pooled subclass level. The VLMC analysis unit represented the source VLMC & Perivascular subclass, comprising VLMC_1, Pericyte_1, Pericyte_2-SEAAD and SMC-SEAAD. Marker displays supported annotation only and were not used for differential-expression inference. Figure 2c displays 48 markers across 24 subclasses; dot size denotes the detected-expression fraction and color denotes gene-scaled mean expression.

#### Donor-level cell proportions and continuous CPS association

For each early-window donor, the seven non-neuronal populations were expressed as percentages of all atlas nuclei from that donor, retaining zero-count combinations. Low and High groups were defined by the early-window median CPS (15 donors each) and compared by two-sided Wilcoxon rank-sum tests, with BH correction across the seven populations (Fig. 2d). Micro-PVM percentage was tested against continuous CPS by two-sided Spearman correlation; a donor-bootstrap percentile 95% interval used 4,000 resamples (seed 20260919; Fig. 2e). Unadjusted and age-, sex- and APOE ε4-adjusted linear fits were descriptive sensitivity analyses.

#### Differential expression analysis

Within each analysis unit and CPS window, we modeled nucleus-level counts using NEBULA, a negative-binomial generalized linear mixed model with a donor-specific random intercept[29]. CPS was fitted as a continuous predictor, with sex, age at death, and *APOE* ε4 carrier status included as covariates. The donor random intercept accounted for within-donor correlation among nuclei. Across the seven principal analysis units, models required at least 20 contributing donors, genes had to be detected in at least five nuclei, and the retained-gene count total for each nucleus was supplied as its library-size offset. The number of contributing donors and nuclei was retained for every model, and model failures or insufficiently detected units were recorded rather than imputed. Effective sample sizes, tested-gene counts and recorded convergence codes are provided in Supplementary Table S3a (Additional file 3).

For each gene, the CPS coefficient, standard error, Wald z statistic, nominal P value, and Benjamini–Hochberg FDR were exported. Multiple-testing correction was performed separately within each analysis-unit-by-window model, and FDR < 0.05 was considered statistically significant. CPS was entered without rescaling; the natural-log coefficient β was exported as log2FC = β/ln(2), so the effect per 0.1 CPS was 0.1 × β/ln(2), equivalently, 0.1 × the exported log2FC. Differentially expressed gene counts were not compared as a measure of relative cell-type responsiveness because cell yield, donor coverage, and gene detection differed substantially among units.

#### Gene set enrichment analysis

Genes were ranked by the signed NEBULA Wald statistic (z = coefficient/standard error) without applying a gene-level significance threshold. Gene set enrichment analysis was performed with fgseaMultilevel using locally stored Molecular Signatures Database (MSigDB) v2026.1 collections: Hallmark, KEGG Legacy, Reactome, Gene Ontology Biological Process (GO BP), and transcription-factor target sets from TFT GTRD[21–24,30]. Gene sets containing 10-500 ranked genes were tested with eps = 0. Across the seven principal analysis units, Benjamini–Hochberg correction was applied jointly across eligible Hallmark, GO BP, Reactome and KEGG Legacy gene sets within each analysis-unit-by-window comparison; TFT GTRD was tested and corrected separately. Original source FDR values were retained for all displayed comparisons. Figure 2f displays 10 inflammation/immunity pathways and six ECM, adhesion and fibrosis pathways. Each displayed pathway reached FDR < 0.05 in at least one analysis unit in the early window. GO BP redundancy was reduced using semantic similarity[25]. Supplementary Fig. S3a displays 34 late-window pathways across five functional themes. Display filtering did not change the source NES or FDR values.

#### Cell-cell communication analysis

Within each CPS window, donors were divided at the window-specific median into low- and high-CPS groups (15 vs 15 donors in the early window; 24 vs 24 donors in the late window). Nuclei were pooled within each group and balanced by downsampling to a maximum of 1,000 nuclei per cell type (seed = 42). Cell types represented by fewer than 10 nuclei were removed and recorded. Low- and high-CPS networks were then estimated separately with CellChat v2 and CellChatDB.human, using computeCommunProb with the trimean estimator and 100 bootstrap iterations, filterCommunication (min.cells = 10), computeCommunProbPathway, and aggregateNet[26,27].

The two networks within each window were merged for descriptive comparison. Figure 2g reports the early-window high-minus-low difference in modeled communication strength for each sender–receiver pair; the corresponding late-window network is shown in Supplementary Figure S3b. For Figure 2h and Supplementary Figure S3c, pathway information flow was normalized as high/(low + high). For pathways detected in both groups, matched ligand–receptor records (sender × receiver × interaction) were compared using a two-sided paired Wilcoxon signed-rank test with a normal approximation and continuity correction (exact = FALSE, correct = TRUE). Missing record-level probabilities were set to zero. Benjamini–Hochberg correction was applied separately across the 96 early-window and 95 late-window pathways with actual test results, before presentation filtering. FDR < 0.05 defined the record-level threshold. Figure 2h was restricted to 42 pathways in the two main-figure classes (25 inflammation/immunity and 17 ECM/adhesion pathways), whereas Supplementary Figure S3c displayed all 102 late-window pathways across five annotated functional classes. Pathways detected in only one group were summarized descriptively (three in the early window and seven in the late window), with P values and FDR recorded as not applicable. They were excluded from the statistical testing family and significant-pathway counts and marked separately in the figures. The TNF communication pathway was absent under the applied expression and detection criteria and was not evaluated as a candidate pathway.

CellChat comparisons were based on pooled, downsampled nuclei. The ligand–receptor record, not the donor, was the unit of the pathway-level significance test. These analyses are therefore descriptive model-based inferences and were interpreted in terms of direction and pathway theme; they do not constitute donor-level confirmation of altered intercellular communication.

#### Statistical reporting

Donors were treated as the biological units for cohort definition and covariate assignment. All tests were two-sided unless intrinsically directional, and multiple-testing correction used the Benjamini–Hochberg procedure[13]. Exact donor and nucleus counts, effect directions, effect statistics, and FDR values are reported where relevant. UMAP was used only for visualization and not for statistical inference.

### Cross-tissue inflammatory convergence in endothelium and myeloid cells (Figures 3–4; Supplementary Figures S4–S5)

#### Input results and analysis populations

Endothelial comparisons used HLMA donor-aggregate limma-GSEA and early-window SEA-AD Endothelial NEBULA-GSEA results. Myeloid comparisons used a separate muscle single-nucleus mixed-effects MAST analysis and early-window SEA-AD Micro-PVM NEBULA-GSEA results. Endothelial heatmaps comprised 23,227 muscle endothelial cells from 23 donors and 693 brain endothelial nuclei from 30 donors. The myeloid comparison retained 4 Young donors (259 nuclei) and 9 Old donors (4,166 nuclei) in muscle and 13,416 brain Micro-PVM nuclei from 30 donors. Muscle exposure was age group and brain exposure was continuous CPS.

#### Inflammatory pathway alignment and selection

Both analyses used the same fixed module of eight Hallmark inflammation/immunity pathways: allograft rejection, complement, inflammatory response, interferon-α response, interferon-γ response, IL-2/STAT5 signaling, IL-6/JAK-STAT3 signaling and TNF-α/NF-κB signaling. Panel a included pathways reaching source FDR < 0.05 separately in each tissue. The shared panel-b set additionally required the same NES direction and was ranked by the larger of the two tissue-specific FDR values, then mean FDR and pathway identifier. Up to six pathways were displayed; six qualified for endothelium and four for myeloid cells. Source FDR values and their original testing families were retained without post-selection recalculation.

#### Leading-edge intersections and functional identities

For each selected pathway, complete native GSEA leading-edge or core-enrichment lists were intersected without top-gene truncation or supplementation with other pathway members[19]. The unions of these pathway-specific cross-tissue intersections contained 109 unique endothelial genes across six pathways and 51 unique myeloid genes across four pathways. Intersections were descriptive; no overlap significance test was applied, and Venn diagrams were not area proportional.

TF identities were assigned from the HumanTFs reference list, and ligand and receptor identities from the local human CellChatDB protein-interaction categories[26,27,31]. Roles were nonexclusive. Main heatmaps displayed the 41 endothelial and 21 myeloid genes with at least one TF, ligand or receptor identity; complete 109- and 51-gene heatmaps were retained in Supplementary Figures S4 and S5. Within each tissue, donor aggregates were scaled per million using totals over all features, log1p-transformed and standardized by gene. Muscle aggregates comprised back-transformed normalized expression, whereas brain aggregates comprised raw counts. Muscle donors were ordered by age and brain donors by CPS, without cross-tissue normalization or missing-gene imputation.

#### Muscle myeloid mixed-effects model and sensitivity analyses

The muscle myeloid reanalysis was restricted to Chinese single-nucleus donors with at least 20 myeloid nuclei. Log2(1 + normalized expression) was modeled in MAST with age group, sex, standardized cellular detection rate and a donor random intercept[32]. The cellular detection rate was the proportion of source features detected in each nucleus. A joint likelihood-ratio test combined the hurdle components. Adjusted mean differences averaged predictions over a common sex and detection-rate distribution with equal donor weights and the random intercept set to zero. Genes were ranked by the sign of the adjusted Old-versus-Young predicted mean difference multiplied by the square root of the joint statistic, and Hallmark GSEA used the complete eligible Hallmark testing family with FDR < 0.05.

Donor-level limma-GSEA and CAMERA were retained as sensitivity analyses[33]. Supplementary Fig. S1 used the same MAST ranking but displayed the inflammation/immunity and ECM, adhesion and fibrosis modules from Fig. 1e at the predefined nominal P < 0.05 threshold. This supplementary display threshold was not used to select the shared Figure 4 pathways. Donor pathway-score tests were also retained as a separate summary and corrected across the tested Hallmark pathways. Model diagnostics, matched-gene-universe comparisons and analyses excluding random-effect boundary fits are detailed in Supplementary Methods and Supplementary Table S4 (Additional file 3).

Shared enrichment required tissue-specific FDR < 0.05 and concordant NES direction; it was not a pooled effect or joint P value. Effect magnitudes from the different tissue models were not directly compared. Muscle myeloid cells and brain Micro-PVM were treated as functionally corresponding populations, not as equivalent lineages or evidence of migration.

### Peripheral-blood single-cell and bulk-transcriptome analyses (Figure 5; Supplementary Figures S6–S8)

#### Study design, datasets, and terminology

The primary analysis examined two public PBMC single-cell RNA-sequencing resources separately. The AD-spectrum analysis used SRP312418 and was restricted to HC, MCI, and AD[10]. After repeat samples were aggregated within diagnosis and two participants represented in both MCI and AD were excluded, participant-level analyses included 121 HC, 46 MCI, and 27 AD participants. Sex was included as a covariate; age was unavailable in the released analysis object, and *APOE* information was too incomplete for the primary model. GSE157007 comprised six healthy older controls and five participants with the source study’s frailty phenotype[34]. The group is labeled Frailty throughout; no clinical sarcopenia diagnosis was inferred from this phenotype. Complementary bulk analyses used GSE206762 leukocyte RNA sequencing and ADNI whole-blood microarray data.

#### UMAP visualization and cell-type harmonization

Fig. 5a displays all 76,509 GSE157007 cells using the existing Harmony-derived embedding[35]. Published SRP312418 UMAP coordinates were joined one-to-one between annotation endpoints for visualization only, and Fig. 5b displays a deterministic diagnosis- and cell-type-stratified sample of 250,096 cells from the 696,093-cell HC/MCI/AD atlas. Display sampling and coordinate matching did not determine participant-level composition or pseudobulk statistics. Detailed embedding and harmonization procedures are provided in Supplementary Methods.

#### Participant-level compositional analysis

Cell-type proportions were calculated using all quality-controlled PBMCs from each participant as the denominator. Mean percentage-point differences from HC and nonparametric bootstrap percentile 95% CIs were estimated by resampling participants within diagnosis groups (2,000 replicates for SRP312418; 10,000 for GSE157007). CIs describe uncertainty in unadjusted mean differences and were not used as compositional significance tests. Primary compositional inference used scCODA[36]. The SRP312418 model included diagnosis and sex, used plasma cells as the reference, and used 1,000 warm-up and 2,000 posterior samples at an estimated FDR of 5%. The GSE157007 model included group, standardized age and source batch, used CD4+ T cells as the reference, and used 1,000 warm-up and 5,000 posterior samples at an estimated FDR of 5%. Figure 5c combines the three contrasts across eight displayed cell categories; black outlines identify effects retained by the respective posterior-selection rule. The NKT-like/T-NK* row aligns broad source labels only and does not establish identical subpopulations.

#### PBMC pseudobulk differential expression and GSEA

Expression matrices were aggregated by participant and broad cell type, retaining the participant as the biological replicate. The released SRP312418 H5AD X matrix contained non-negative fractional counts on the linear expression scale. Differential expression was analyzed with limma-voom[18,37]. Each participant–cell-type pseudobulk required at least 20 cells, with at least 10 retained participants per group. Genes were filtered using edgeR::filterByExpr with the design matrix, min.count = 10 and min.total.count = 15. TMM normalization was followed by voom precision weighting and robust empirical Bayes moderation with a diagnosis-plus-sex design; genes were ranked by the moderated t statistic[17]. Effective HC/MCI/AD sample sizes for the six displayed cell types were 121/46/27 for CD4+ T, 120/45/27 for CD8+ T, 115/42/26 for NK, 114/43/24 for B cells, 93/38/19 for monocytes and 77/29/16 for NKT-like cells. GSE157007 used integer-verified RNA/counts raw counts, requiring at least 10 cells per pseudobulk and three participants per group; all six displayed cell types retained six HC and five Frailty participants. After design-aware expression filtering with edgeR::filterByExpr (min.count = 10) and TMM normalization, edgeR robust quasi-likelihood negative-binomial models included source batch, group and standardized age[16,38]. Genes were ranked by sign(log2 fold change) multiplied by the square root of the quasi-likelihood F statistic. Each disease or Frailty group was compared with HC from its own cohort. Gene-level Benjamini-Hochberg correction was applied separately across tested genes within each cell-type-by-contrast analysis.

The main-figure pathway analysis used the frozen MSigDB 2026.1.Hs Hallmark collection and fgseaMultilevel, retaining gene sets with 15–500 ranked genes[21]. Rankings were the moderated t statistic for SRP312418 and sign(log2 fold change) multiplied by the square root of the quasi-likelihood F statistic for GSE157007. Antibody-derived features were excluded from enrichment rankings. For each contrast, Benjamini–Hochberg correction was applied across all eligible Hallmark pathway-by-cell-type combinations in the six main PBMC populations as one prespecified family. A positive normalized enrichment score denotes enrichment in MCI, AD or Frailty relative to the corresponding cohort-specific HC group.

#### Source-related PBMC pathway and gene displays

Figure 5d used six Hallmark pathways defined by the tissue intersections: IL-6/JAK-STAT3 signaling, interferon-γ response, inflammatory response, TNF-α/NF-κB signaling, allograft rejection and complement. Each was displayed across monocytes, CD4+ T, CD8+ T, B, NK and NKT-like cells in Frailty versus HC, MCI versus HC and AD versus HC. The same pathway and cell-type order was retained across contrasts, including nonsignificant estimates. The display was a fixed biological subset of the complete corrected Hallmark testing family and was not a separate six-pathway multiple-testing analysis.

Figures 5e and 5f projected the complete 109-gene endothelial and 51-gene myeloid tissue intersections into the PBMC differential-expression results. A gene was retained for display when log2 fold change > 0 and unadjusted P < 0.05 in at least one cell-type–contrast context; all available tests for retained genes were then represented using the stated display encoding. Gene-level FDR values remained those from the complete original differential-expression testing family and were not recalculated over the displayed subset. Pale entries denote tested results that did not meet the positive nominal display rule, whereas crosses denote unavailable tests. A gene could recur across cell types or contrasts; reported contrast-level counts refer to unique genes and do not represent independent tests.

#### GSE206762 peripheral-blood leukocyte bulk RNA sequencing

GSE206762 assayed bulk RNA from red-blood-cell-lysed peripheral-blood leukocytes, not PBMCs or PAXgene whole blood[39]. The source cohort comprised 35 frail and 35 robust participants older than 70 years. Public RNA sequencing was performed on 20 sex-stratified RNA pools: 10 pools per phenotype, including five female and five male pools in each group. The 20 RNA pools were the statistical replicates; the 70 contributing participants were not analyzed as independent RNA-sequencing libraries. The source frailty phenotype required a timed up-and-go of at least 12 s, gait speed of 0.8 m/s or lower, and a Short Physical Performance Battery score below 10; Robust controls met the reciprocal criteria. The bulk figures retain the source label Robust.

Counts were filtered and TMM normalized in edgeR[16,17]. Frailty was compared with Robust using a robust quasi-likelihood generalized linear model with sex and condition as covariates[38]. Pool-level age was unavailable and was not included. Genes were ranked by sign(log2 fold change) multiplied by the square root of the quasi-likelihood *F* statistic. Multilevel GSEA used Hallmark, Reactome, KEGG Medicus, and WikiPathways gene sets containing 10-500 genes[21,22,30,40]. Among 19,389 genes tested, 1,654 were nominally increased and 1,429 nominally decreased at *P* < 0.05, but none reached a gene-level BH-adjusted FDR below 0.05; gene-level findings were therefore considered exploratory.

#### ADNI whole-blood analysis and cross-modality comparison

An independent AD-spectrum system-level comparison used ADNI PAXgene whole-blood microarray data from 244 HC, 376 MCI, and 112 AD participants[41]. MCI and AD were each compared with the common HC group using limma with robust, intensity-trend empirical Bayes moderation and adjustment for age, sex, RNA integrity number, study phase, and Affymetrix array plate[18]. Genes were ranked by the moderated *t* statistic for GSEA. Additional cohort and access information is provided in the Supplementary Methods.

For Supplementary Fig. S6, fixed Fig. 3 endothelial and Fig. 4 myeloid intersection-gene subsets were defined separately within the six source-related Hallmark pathways and tested against the complete eligible bulk rankings. fgseaMultilevel used source-subset sizes of 5–500, with sampleSize = 1,001, nPermSimple = 10,000, eps = 0 and gseaParam = 1. Benjamini–Hochberg correction was applied across the 30 prespecified source-by-pathway-by-comparison tests. Fig. 4 supplied no Complement or TNF-α/NF-κB subset, which was coded as undefined rather than nonsignificant. Supplementary Fig. S7 retained endothelial-source genes and Supplementary Fig. S8 retained myeloid-source genes with log2 fold change > 0 and unadjusted P < 0.05 in at least one bulk comparison; both displayed all available effects across GSE206762 Frail versus Robust, ADNI MCI versus HC and ADNI AD versus HC. Gene-level FDR values remained those from each complete original testing family.

#### Statistical reporting and mechanistic interpretation

Frequentist tests were two-sided unless direction was inherent to the enrichment procedure. PBMC pathway significance used Benjamini–Hochberg FDR < 0.05 across all eligible Hallmark pathway-by-cell-type tests within each contrast. Supplementary Fig. S6 used FDR < 0.05 across its prespecified 30-test family, whereas PBMC and bulk gene panels retained FDR values from the complete original gene-level testing backgrounds. Composition shifts with bootstrap CIs excluding zero but without scCODA posterior credibility were interpreted descriptively. Significance in one comparison but not another was not treated as a test of differences between effects. Endothelial-source and myeloid-source labels indicate provenance from tissue intersections and do not assign a circulating cell of origin in bulk blood. The cross-sectional PBMC and bulk cohorts do not constitute a longitudinal Frailty-to-MCI-to-AD sequence, and clinical MCI is not equivalent to the SEA-AD early CPS window.

### Cluster-specific CellOracle in silico perturbation analyses (Figure 6; Supplementary Figures S9–S10)

#### Candidate factors, analysis populations and estimability

The Fig. 3 intersection supplied 18 endothelial TF candidates, and the Fig. 4 intersection supplied six myeloid candidates (IRF1, STAT3, SPI1, STAT2, MXD1 and PLSCR1). All candidates were retained in the network-support displays. The pre-fit screen required TF detection in at least 10 profiles and 1% of profiles, at least three donors with three or more expressing profiles each, and at least five promoter-prior targets among the selected features. A tissue-level knockout was considered estimable when the TF passed this expression and promoter-prior screen and at least one eligible state contained five or more fitted outgoing TF–target edges. One to four fitted edges were considered insufficient support, and zero fitted edges were recorded separately. Insufficient expression, absent promoter-prior support or inadequate fitted outdegree was coded as not estimable rather than no biological effect. Fifteen endothelial candidates were estimable in at least one tissue and 13 in both; four myeloid candidates were estimable in both tissues.

#### State definitions, features and cluster-specific regulatory models

The endothelial analysis used 2,753 HLMA profiles from 15 Old donors across four eligible states and 640 SEA-AD nuclei from 30 early-window donors (CPS ≤ 0.55; Severely_Affected = N) across two states. Muscle states were displayed as Capillary-like, Stress response, Venous-like and Remodeling-like from marker evidence; these labels were descriptive or provisional, and the Stress response state was strongly enriched for scRNA profiles. Brain Endo_2 and Endo_3 were displayed as BBB capillary-like and inflammatory venular-like, respectively, while retaining the source state identities. A small muscle state was excluded because its muscle-lineage marker program had a mean detection fraction of at least 0.6 and mean log1p expression more than twice that of the endothelial marker program; this did not establish that its cells were doublets. The supplementary myeloid analysis used 471 HLMA snRNA profiles from eight Old Chinese donors across two marker-informed provisional states (Activation-like and Glycolytic inflammatory-like) and 8,200 SEA-AD Micro-PVM nuclei from 30 early-window donors across five retained source states. Each donor contributed at most 300 randomly sampled profiles (seed 20260921). Eligible states contained at least 100 profiles and at least five donors contributing three or more profiles each. Harmony-corrected coordinates were used only for muscle endothelial state display; regulatory models used unintegrated expression, and an snRNA-only endothelial reconstruction was retained as a sensitivity analysis.

For each lineage, shared genes detected in at least 10 profiles and 1% of profiles in each tissue were ranked by the mean of their tissue-specific normalized-dispersion percentile ranks. The top 3,000 features were fixed while retaining eligible candidate TFs. CellOracle 0.20.0 used a public hg38 promoter Base-GRN without matched cell-type-specific chromatin-accessibility data[11]. Normalized log1p expression was imported into CellOracle and smoothed using balanced k-nearest-neighbor imputation over 30 principal components, with k set to 2.5% of profiles, rounded to the nearest integer, and a minimum of 10. State-specific networks were fitted with ridge α = 10 and 20 rounds of cell bagging. Edges were filtered at model P ≤ 0.001 and by absolute coefficient, with at most 2,000 edges retained per state. For each estimable TF, expression was set to zero and the predicted changes were propagated for three steps. These simulations predicted expression changes in the fitted feature space and did not represent experimental knockout cells.

#### Program responses, enrichment and candidate prioritization

For each lineage, tissue and TF, predicted knockout-minus-wild-type changes were averaged equally across retained profiles within each donor and then averaged with equal donor weights. For an estimable TF, predictions from every retained state contributed, including states with fewer than five fitted outgoing edges; no state was discarded or assigned a placeholder zero on this basis. All 15 muscle endothelial, 30 brain endothelial, eight muscle myeloid and 30 brain myeloid donors contributed, with no missing donor predictions. Source programs used the full Hallmark memberships of the six Fig. 3 endothelial inflammatory programs or four Fig. 4 myeloid programs, restricted to modeled genes. Scores excluded the directly zeroed TF, required at least 10 remaining members and averaged gene-wise standardized donor responses. Standardization used each gene’s imputed wild-type expression SD across all retained profiles in that tissue and lineage, floored at 0.1 (Supplementary Methods). Conditional 95% CIs used 2,000 donor bootstrap resamples. Exact two-sided donor sign tests were Benjamini–Hochberg-corrected across the corresponding source programs within each knockout. The program score is a custom predicted-response summary, not a CellOracle trajectory perturbation score. Because donor predictions shared fitted networks, these intervals and tests describe conditional model output and do not capture uncertainty from network estimation.

GSEA ranked donor-equal predicted Δlog1p values after excluding the perturbed TF and used the frozen Hallmark 2026.1.Hs collection. Global Benjamini–Hochberg correction was applied within each tissue and lineage across all estimable TFs and eligible Hallmark tests. The cross-tissue evidence audit recorded the signs of program scores and conditional intervals, GSEA direction and global FDR, five mutually exclusive donor-group-out network rebuilds, and six parameter variants; endothelial evidence also required concordance in the muscle snRNA-only reconstruction. A fully supported TF–program pair required negative responses and intervals in both tissues, global GSEA FDR < 0.05 in both, and concordance across all applicable robustness checks. No endothelial or myeloid pair met this rule. ETS2 and BHLHE40 were prioritized as exploratory endothelial candidates because each had the largest number of programs meeting the exploratory criteria; this ranking did not establish pathway inhibition after multiple-testing correction. The exploratory criteria and candidate ordering are defined in Supplementary Methods, with the complete program-level audit and endothelial ranking in Supplementary Table S5 (Additional file 3).

#### State-space vectors and randomized-residual controls

For every TF estimable in at least one tissue, CellOracle transition probabilities were projected onto the observed two-dimensional state spaces. The main figure displays ETS2 and BHLHE40. Supplementary Fig. S9d displays all 18 endothelial candidates, with vector maps for the 15 candidates estimable in at least one tissue and gray placeholders for PLSCR1, RBCK1 and TRAFD1, which were not estimable in either tissue. Supplementary Fig. S10e similarly displays the four estimable myeloid candidates and gray placeholders for MXD1 and PLSCR1. Placement within these atlas panels has no prescribed order and does not indicate rank. A native randomized-residual calculation was processed through the same projection pipeline as a descriptive control. Within each tissue, knockout and randomized maps used identical grid points, thinning, arrow scale, color, width and opacity; scales were not compared across independent tissue embeddings. Grid points were not treated as biological replicates, and a single randomized field did not provide a biological P value or null distribution. No pseudotime root or Young-state trajectory was imposed.

#### Core-program and leading-edge gene summaries

Figure 6e retained three selected negative program responses for each prioritized TF: interferon-γ response, inflammatory response and complement for ETS2; and IL-6/JAK-STAT3 signaling, interferon-γ response and TNF-α/NF-κB signaling for BHLHE40. Effects and bootstrap intervals were donor-equal program scores; exact sign-test P values and within-knockout Benjamini–Hochberg q values were reported separately from global GSEA FDR.

#### Figure 6f gene selection

Figure 6f selected genes that belonged to the same TF–program GSEA leading edge in both tissues, had negative predicted effects in both, and passed gene-level Benjamini–Hochberg FDR < 0.05 in both tissue models. Gene FDR was based on donor sign tests across the modeled feature background within each knockout. Direct filtered-network targets and propagated responses were distinguished in source data; neither category established experimental binding or causal regulation.

Five mutually exclusive donor-group-out rebuilds repeated imputation, bagged network inference, edge filtering and propagation with fixed features and state labels. Additional sensitivity analyses varied retained edge counts, propagation steps, knockdown strength and post-propagation within-state range constraints; muscle endothelium also used an snRNA-only reconstruction. Rebuilds used their own baseline standard deviations and were interpreted as robustness checks rather than independent biological replicates. All effects remained conditional on the observational expression data, promoter prior, selected features and fitted CellOracle models. Cross-tissue agreement therefore supports candidate prioritization only and does not establish pathway inhibition in vivo, endothelial or myeloid state rescue, or causal muscle-to-brain signaling.

## RESULTS

### Inflammatory and matrix remodeling in aging muscle accompanied stromal-dominant inferred communication

To establish a muscle-side reference for age-associated molecular changes, we analyzed a public human limb skeletal muscle atlas (HLMA) containing single-cell RNA-sequencing and single-nucleus RNA-sequencing (snRNA-seq) data from 292,423 cells and nuclei across 23 donors[4]. These comprised 8 Young donors aged 15–45 years and 15 Old donors aged 77–99 years. The atlas resolved 15 populations, including myonuclei, muscle stem/progenitor cells, stromal cells, vascular cells, immune cells, adipocytes, Schwann cells and erythroid cells (**Fig. 1a**). Population-restricted expression across a 30-gene marker panel supported the original annotations (**Fig. 1b**), and both age groups contained the major populations.

In the shared snRNA-seq subset, none of the 15 populations differed in donor-level proportion at a false discovery rate (FDR) below 0.05 after multiple-testing correction. **Fig. 1c** displays nine muscle, stromal and vascular populations. Myeloid-cell and lymphocyte proportions were descriptively higher in Old but likewise did not reach the adjusted threshold (**Fig. 1d**). We therefore focused on transcriptional programs within cell types to characterize age-associated changes.

Inflammation-related pathway enrichment was distributed across the 11 evaluable cell types, including the three lower-powered populations analyzed separately. TNF-α/NF-κB signaling was positively enriched in nine, accompanied by increased interferon-γ response, complement, IL-6/JAK-STAT3 signaling and cytokine–receptor interaction. Stromal and vascular-wall populations also showed extracellular-matrix (ECM)/adhesion remodeling. Epithelial–mesenchymal transition was positively enriched in 8 of 11 cell types, with increased TGF-β signaling and ECM organization, alongside reduced focal adhesion in fibro/adipogenic progenitors (FAPs), tight-junction signaling in endothelial cells and ECM–receptor interaction in FAPs and lymphocytes (**Fig. 1e**). These findings identified broad inflammatory and vascular–stromal transcriptional remodeling across aging muscle cell populations.

These transcriptional changes accompanied altered patterns of inferred intercellular communication. Relative to Young, the pooled Old network contained 77% more ligand–receptor (L-R) interactions and approximately 75% greater total modeled communication strength. FAPs and fibroblast-like cells showed the largest increases in outgoing signaling, with myonuclei, muscle satellite cells (MuSCs), endothelial cells and other populations among the recipients (**Fig. 1f**). This pooled model identified stromal populations as prominent senders of age-associated communication changes, without providing a donor-level test of communication effects.

At the pathway level, ANNEXIN, CypA, GALECTIN and SPP1 were detected only in the pooled Old network, whereas relative information flow for CXCL, COMPLEMENT, MIF, COLLAGEN, LAMININ, THBS and FN1 was skewed toward Old (**Fig. 1g**). This inflammatory and matrix-remodeling profile provided the muscle-side reference for asking whether corresponding vascular–immune programs were present in AD-related brain tissue.

### The early CPS window showed vascular–immune transcriptional activation and redistribution of inferred communication

To assess whether AD-related brain tissue contained molecular programs corresponding to those in aging muscle, we analyzed 1,178,694 nuclei from 84 donors aged 65–102 years in the SEA-AD middle temporal gyrus atlas[28]. Cell-type and donor-level CPS maps displayed the major neuronal and non-neuronal compartments (**Fig. 2a,b**). A 48-gene marker panel across 24 subclasses supported the retained annotations (**Fig. 2c**). Subsequent analyses focused on seven non-neuronal populations: Micro-PVM, Lymphocyte, Astrocyte, Oligodendrocyte, oligodendrocyte precursor cell (OPC), Endothelial and vascular and leptomeningeal cell (VLMC). The differential-expression and enrichment unit labeled VLMC pooled the VLMC & Perivascular subclass. Immune marker expression further supported the distinction between Micro-PVM and Lymphocyte (Supplementary **Fig. S2a,b**).

We estimated local associations within two intentionally overlapping CPS windows to compare transcriptional and communication patterns across ranges of pathological burden. The early window comprised 30 donors with CPS ≤ 0.55; the late window comprised 48 donors with CPS > 0.45. Five donors contributed to both windows. The main figure focused on the early window, with the late window retained as a contrast in Supplementary **Fig. S3**. Both were cross-sectional analytical ranges, not longitudinal disease stages in the same donors.

Donor-level proportions of the seven non-neuronal populations did not differ between the Low-and High-CPS groups after FDR correction (**Fig. 2d**). No monotonic association was detected between Micro-PVM abundance and continuous CPS (Spearman ρ = 0.004, P = 0.982; **Fig. 2e**). These tests did not detect composition associations, but do not exclude smaller changes or shifts within the pooled populations.

Within cell types, gene set enrichment analysis identified inflammation/immunity and ECM, adhesion and fibrosis programs at the vascular–immune interface (**Fig. 2f**). Interferon-γ responses were positively enriched in Micro-PVM, Lymphocyte, Endothelial and VLMC populations. IL-6/JAK-STAT3 enrichment involved Micro-PVM, Endothelial and VLMC, whereas TNF-α/NF-κB enrichment was significant in Lymphocyte, Endothelial and VLMC. Adhesion and junctional responses were cell-type specific, combining positive cell–cell adhesion regulation with negative enrichment of selected junctional programs.

To examine accompanying changes in inferred cell–cell communication, we compared CellChat networks between the low- and high-CPS groups. Inferred interactions decreased from 2,964 to 2,786 between low- and high-CPS groups, whereas total modeled communication strength remained similar (111.36 versus 112.52). Sender–receiver changes varied in direction across cellular compartments (**Fig. 2g**). Higher CPS within the early window was therefore associated with redistribution of modeled signaling rather than global amplification of the communication network.

Pathway-level analysis revealed a directional shift within this otherwise stable global network. Among the 42 displayed inflammation/immunity and extracellular-matrix (ECM)/adhesion pathways, 11 met the ligand–receptor-record-level false discovery rate threshold, and all showed greater relative information flow in the high-CPS group (**Fig. 2h**). These included SPP1 and transforming growth factor-β (TGF-β) signaling together with the matrix-associated COLLAGEN, LAMININ, FN1, and THBS pathways. ANNEXIN and CLEC were detected only in the high-CPS network and were summarized descriptively, separately from the tested differences. Thus, early-window inferred communication was selectively redistributed toward inflammatory and ECM/adhesion programs.

By contrast, the late CPS window showed more heterogeneous and cell-type-specific transcriptional responses (Supplementary **Fig. S3a**). Total modeled communication strength was 19.7% lower in the high-CPS group, and 38 of 39 pathways meeting the record-level FDR threshold showed lower relative information flow, indicating broad attenuation of inferred communication rather than persistence of the early-window pattern (Supplementary **Fig. S3b,c**).

The early-window vascular–immune pattern, contrasted with broader communication attenuation in the late window, motivated its selection for subsequent muscle–brain comparisons of shared cellular programs.

### Muscle and brain endothelial cells shared inflammatory pathways and molecular components

Having characterized both tissues independently, we examined whether endothelial inflammation involved the same pathways. Within a fixed module of eight Hallmark inflammation/immunity pathways, seven were significant in each tissue and six were shared at FDR < 0.05 with concordant positive enrichment toward Old in muscle and higher CPS in brain (**Fig. 3a,b**). These comprised IL-6/JAK-STAT3 signaling, interferon-γ response, inflammatory response, TNF-α/NF-κB signaling, allograft rejection and complement.

Complete leading-edge intersections across the six pathways yielded 109 unique genes (**Fig. 3c**). Of these, 41 had transcription-factor (TF), ligand or receptor identities (Fig. 3d), including STAT1, STAT3, NFKB1, RELA, IL6, CCL2, IL6ST, IL1R1 and IFNAR1. BHLHE40, JUNB and IRF2 were also present. These annotations identify candidate molecular roles, not activity or signaling direction.

Heatmaps displayed the 41 role-annotated genes across 23 muscle and 30 brain donors, ordered by age and CPS, respectively (**Fig. 3e**). The complete 109-gene view was retained in Supplementary **Fig. S4**. Together, these analyses defined a shared endothelial inflammatory program and motivated comparison of the myeloid compartments.

### Muscle myeloid cells and brain Micro-PVM shared inflammatory pathways and genes

We next asked whether inflammatory convergence extended beyond endothelium. Because Young muscle donors contributed relatively few myeloid nuclei, we used a targeted MAST analysis with a donor random effect in the eligible single-nucleus subset (4 Young and 9 Old donors) and compared it with Micro-PVM results from 30 early-window brain donors. Four pathways were positively enriched in both tissues at FDR < 0.05: inflammatory response, allograft rejection, interferon-γ response and IL-6/JAK-STAT3 signaling (**Fig. 4a,b**).

Intersecting the complete leading-edge sets yielded 51 unique shared genes (**Fig. 4c**), of which 21 had TF, ligand or receptor identities (**Fig. 4d**). These included IRF1, STAT3, CCL2, TGFB1, IFNAR1 and IL10RA. Donor-level heatmaps showed the 21 annotated genes across muscle and brain (**Fig. 4e**); the complete 51-gene set was retained in Supplementary **Fig. S5**.

The supplementary muscle MAST analysis also examined the inflammation/immunity and ECM, adhesion and fibrosis modules used in **Fig. 1e**. Nine of the 16 displayed pathways met the predefined nominal P < 0.05 display threshold (Supplementary **Fig. S1**). Donor-level limma-GSEA supported the positive direction of the four shared inflammatory pathways, whereas donor pathway-score tests did not survive multiple-testing correction. The myeloid result therefore represents cross-tissue functional correspondence, not equivalence or migration between muscle myeloid cells and brain Micro-PVM.

### Tissue-derived inflammatory programs showed context-dependent PBMC patterns and external support in bulk blood

After identifying shared inflammatory programs in aging muscle and AD-related brain tissue, we examined their systemic immune correlates in independent peripheral-blood cohorts.

GSE157007 comprised 76,509 peripheral blood mononuclear cells (PBMCs) in 11 cell types from six healthy older controls and five participants with source-defined Frailty (**Fig. 5a**)[34]. The SRP312418 atlas comprised 696,093 PBMCs from cognitively healthy controls (HC), participants with mild cognitive impairment (MCI), and participants with AD, resolved into 12 cell types (**Fig. 5b**)[10]. Frailty denotes the source phenotype and was not treated as clinically diagnosed sarcopenia. Each case group was compared with its own cohort-specific HC group.

Cell-composition changes did not follow a common pattern across the three contrasts (**Fig. 5c**). Monocyte proportions were descriptively higher in MCI and Frailty, but neither met the prespecified scCODA posterior credibility criterion. Frailty also showed descriptively higher NK- and B-cell proportions and lower CD8+ T-cell proportions. Only the increase in AD CD4+ T cells met this criterion, with a mean difference of 5.62 percentage points (bootstrap 95% confidence interval [CI], 0.24 to 10.79). We therefore examined transcriptional programs within cell types to assess whether the cohorts shared functional changes.

We next tested six source-related Hallmark pathways from the endothelial and myeloid tissue intersections across six broad PBMC populations (**Fig. 5d**). After Benjamini–Hochberg correction across all eligible Hallmark pathway-by-cell-type tests within each contrast, 30 of 36 displayed combinations were positively enriched in Frailty, compared with one in MCI and five in AD; no displayed combination showed significant negative enrichment. In Frailty, interferon-γ response, inflammatory response, TNF-α/NF-κB signaling and allograft rejection were significant in all six populations. IL-6/JAK-STAT3 signaling was significant in monocytes, CD4+ T, B and NK cells, and complement in monocytes and B cells. MCI showed only monocyte TNF-α/NF-κB enrichment. AD showed interferon-γ response in CD8+ T and NK cells, TNF-α/NF-κB signaling in CD8+ T cells, and allograft rejection in CD8+ T and B cells.

We then projected the complete 109-gene endothelial and 51-gene myeloid tissue intersections into the PBMC differential-expression results (**Fig. 5e,f**). Genes with log2 fold change > 0 and unadjusted P < 0.05 in at least one displayed cell-type–contrast context were retained, yielding 61 endothelial-source and 32 myeloid-source genes. Across unique genes, Frailty contained 51 endothelial-source and 23 myeloid-source nominally increased genes, compared with 11 and 8 in MCI and 24 and 8 in AD, respectively. Gene-level FDR < 0.05 identified 11 endothelial-source and one myeloid-source gene in Frailty, none in MCI, and nine endothelial-source and four myeloid-source genes in AD. JUNB, SOCS1 and TGIF1 were among the FDR-supported endothelial-source genes in both Frailty and AD; BTG2 was the single supported myeloid-source gene in Frailty, whereas EIF3D, NCF4, SPI1 and TGFB1 were supported in AD.

To assess the tissue-derived programs in independent blood datasets, we tested predefined gene subsets from the Fig. 3 endothelial and Fig. 4 myeloid intersections in the GSE206762 frailty bulk-blood cohort and ADNI MCI/AD whole-blood comparisons. All 30 testable source-by-pathway-by-comparison combinations had positive normalized enrichment scores, and 23 passed Benjamini–Hochberg correction across the prespecified 30-test family (Supplementary **Fig. S6**). For endothelial-source subsets, five of six pathways were supported in GSE206762 Frail versus Robust, four of six in ADNI MCI versus HC, and five of six in ADNI AD versus HC. The corresponding myeloid-source counts were four of four, three of four and two of four. Complement and TNF-α/NF-κB signaling were undefined for the myeloid source because **Fig. 4** yielded no corresponding intersection subset; gray cells therefore did not denote nonsignificance.

Gene-level bulk results were more limited. Supplementary **Fig. S7** displayed complete effect estimates for 38 of 109 endothelial-source genes that were nominally increased in at least one bulk comparison, and Supplementary **Fig. S8** displayed the corresponding 18 of 51 myeloid-source genes. None passed gene-level FDR < 0.05 in any bulk dataset; open circles identify nominal P < 0.05. Recurrent nominal increases across comparisons showed directional agreement but remained exploratory.

The PBMC and bulk-blood analyses therefore supported related inflammatory programs with distinct cohort and cell-type patterns. Frailty had more significant displayed PBMC pathway–cell-type combinations than MCI or AD, whose patterns were cell dependent. These counts were not formal comparisons of effects between cohorts. Because genes could recur across cell types and contrasts, the displayed counts were descriptive rather than independent pieces of evidence. Bulk expression could not assign signals to a circulating cell of origin, and the endothelial-source and myeloid-source labels indicate provenance from the tissue intersections rather than endothelial or myeloid identity in blood. We next used in silico perturbations to prioritize candidate regulators of the shared endothelial programs.

### Cluster-specific perturbation models prioritized ETS2 and BHLHE40 in endothelium but identified no stable myeloid candidate

The **Fig. 3** intersection yielded 18 endothelial TF candidates. CellOracle models included 2,753 muscle endothelial profiles from 15 Old donors across four marker-informed states and 640 early-window brain endothelial nuclei from 30 donors across two retained states (**Fig. 6a**). Fifteen candidates were estimable in at least one tissue, and 13 were estimable in both; unavailable combinations reflected insufficient expression, promoter-prior support or fitted outgoing edges and were not interpreted as evidence of no biological effect (Supplementary **Fig. S9a**).

Across the six source-related endothelial programs, predicted knockout responses were summarized within donors and then averaged with equal donor weights (**Fig. 6b**; Supplementary **Fig. S9b**). No TF–program pair met the prespecified composite support rule, and no source program passed global GSEA FDR < 0.05 in either tissue; the minimum global FDR was 0.9891 in muscle and 0.9777 in brain. The full evidence audit therefore supported candidate prioritization by directional robustness rather than multiplicity-corrected pathway inhibition (Supplementary **Fig. S9c**).

ETS2 and BHLHE40 each had three programs meeting the exploratory prioritization criteria, the highest count among the 13 cross-tissue-estimable TFs. ETS2 was prioritized for interferon-γ response, inflammatory response and complement, whereas BHLHE40 was prioritized for IL-6/JAK-STAT3 signaling, interferon-γ response and TNF-α/NF-κB signaling. For both TFs, the projected knockout vector fields differed descriptively from their matched randomized-residual controls in muscle and brain (ETS2, **Fig. 6c**; BHLHE40, **Fig. 6d**); these maps were not used as statistical tests of perturbation efficacy. Supplementary **Fig. S9d** shows the complete 18-candidate atlas, comprising maps for the 15 candidates estimable in at least one tissue and gray placeholders for the three candidates that were not estimable in either tissue; placement does not denote ranking.

For the six selected TF–program combinations, donor-level predicted effects were negative in every donor in both tissues (**Fig. 6e**). In muscle, the exact two-sided sign-test P value was 6.10 × 10□□ for each program and the within-KO BH q value was 9.16 × 10□□ ; in brain, both P and q were 1.86 × 10□□. These conditional tests summarized donor-level directional consistency of the custom program score and did not supersede the nonsignificant globally corrected GSEA results.

Cross-tissue directional leading-edge filtering identified five predicted downregulated genes for ETS2—CFH, TIMP1, NFKBIA, IL1R1 and DUSP5—and 16 for BHLHE40, including JUN, NAMPT, SOCS3, STAT3, ZFP36, JUNB, SLC2A3 and FOSL2 (**Fig. 6f**). Selection required membership in the same TF–program leading edge in both tissues, negative predicted effects in both, and gene-level FDR < 0.05 in both models. Because the parent GSEA results were not globally significant, these genes remain directional exploratory candidates rather than confirmed pathway targets.

We applied the same framework to six myeloid TFs from the Fig. 4 intersection using 471 muscle myeloid nuclei from eight Old donors and 8,200 early-window brain Micro-PVM nuclei from 30 donors (Supplementary **Fig. S10a**). IRF1, STAT3, SPI1 and STAT2 were estimable, whereas MXD1 and PLSCR1 lacked sufficient network support (Supplementary **Fig. S10b**). No TF–program pair met the stable-negative rule or reached global GSEA FDR < 0.05 in both tissues; the minimum global FDR was 0.9619 in muscle and 0.8709 in brain (Supplementary **Fig. S10c,d**). Supplementary **Fig. S10e** shows maps for the four estimable TFs together with gray placeholders for MXD1 and PLSCR1; the atlas remained descriptive and did not nominate a myeloid TF for the main figure.

## DISCUSSION

In this cross-tissue analysis of independent human transcriptomic datasets, we identified convergent inflammatory programs in endothelial and myeloid populations across aging skeletal muscle and AD-related brain tissue. Their cellular organization nevertheless differed. Aging muscle showed broad inflammatory and extracellular-matrix remodeling across stromal, vascular, immune and myogenic populations, accompanied by stromal-dominant changes in inferred intercellular communication. In SEA-AD middle temporal gyrus, inflammatory and adhesion programs were associated with increasing pathological burden within the early CPS window and concentrated at the vascular–immune interface. Related PBMC signatures and external pathway-level support from the frailty bulk-blood cohort and ADNI MCI/AD comparisons extended these findings to independent peripheral-blood datasets. This convergence is compatible with a putative muscle–brain relationship, but does not demonstrate an interorgan axis or identify a tissue of origin or causal direction.

The muscle findings are consistent with an age-associated tissue state in which inflammatory activation accompanies remodeling of stromal and matrix-related programs[4,5]. Inflammatory enrichment was broadly distributed across cell types, whereas donor-level cell-proportion differences did not survive multiple-testing correction. This combination supports changes within cellular states as a prominent component of aging muscle, although modest composition effects remain possible. FAPs and fibroblast-like cells showed the largest increases in outgoing modeled communication, with myogenic, vascular and other populations among the recipients. Experimental work in mice has likewise linked altered FAP-derived matricellular signaling to impaired muscle stem-cell function during aging[42]. This pattern argues against reducing the muscle-side signal to changes in a single immune population or explaining it solely by cell abundance. Because CellChat comparisons pooled cells within groups, they describe modeled changes in signaling organization rather than donor-level evidence of altered communication.

The brain analysis revealed both convergence and divergence relative to aging muscle. Within the early CPS window, inflammatory and adhesion-related enrichment was most coherent in Micro-PVM, endothelial, VLMC and lymphocyte populations. Selected inflammatory and matrix/adhesion communication pathways showed greater relative information flow despite little change in total modeled communication strength. This combination points to selective reorganization of modeled signaling among cellular compartments, rather than uniform amplification of the network. In the late CPS window, transcriptional responses were more heterogeneous and modeled communication was broadly attenuated. These local analyses describe different cross-sectional patterns. The early CPS window therefore provides a local cross-sectional brain reference for cross-tissue comparison, not a longitudinal ‘early AD’ stage or an equivalent of clinical MCI.

A central finding was that cross-tissue convergence did not imply identical cellular distributions. Endothelial cells offered a comparable compartment in which to examine shared molecular programs without assuming equivalent functions across tissue-specific vascular beds[7,9]. Six concordantly enriched Hallmark pathways and 109 unique shared leading-edge genes defined the endothelial intersection. Shared components included IL6, CCL2, STAT3, IRF2, BHLHE40, JUNB, SOCS1 and SOCS3, placing the overlap within established NF-κB and JAK–STAT biology[43,44]. Role annotation organizes candidates for follow-up but does not establish their activity or signaling direction. The overlap may reflect shared systemic exposures or parallel responses to aging and pathology; it does not distinguish these explanations or establish muscle-derived signaling.

The myeloid comparison extended this convergence beyond endothelium. Four inflammatory pathways and 51 leading-edge genes were shared between muscle myeloid cells and brain Micro-PVM. This correspondence supports a recurring immune program, but the populations are not equivalent and the analysis does not imply migration or a common tissue origin. The mixed-effects muscle analysis addressed donor clustering, while sensitivity analyses did not reproduce every signal. The myeloid component should therefore be interpreted as cross-tissue functional correspondence rather than mechanistic confirmation.

The blood findings extend evidence for systemic immune involvement in AD, while showing that tissue-derived inflammatory programs need not occupy the same cell types across cohorts[45]. TNF-α/NF-κB, interferon-related, inflammatory-response and allograft-rejection programs recurred across PBMC and bulk-blood comparisons, but Frailty had the largest number of significant displayed PBMC pathway–cell-type combinations. MCI and AD showed narrower cell-dependent patterns. Independent bulk-blood analyses of GSE206762 Frail versus Robust and ADNI MCI/AD versus HC provided pathway-level support for the fixed tissue-intersection subsets, whereas gene-level bulk signals remained nominal. These findings are consistent with systemic inflammatory correlates of the muscle–brain relationship, but do not establish blood as a mechanistic bridge between the tissues. The cohorts were independent and unpaired, and Frailty represented the source study’s phenotype rather than sarcopenia diagnosed according to contemporary clinical criteria[46].

The CellOracle analysis prioritized ETS2 and BHLHE40 as exploratory candidate regulators of source-related endothelial programs[11]. Among all endothelial candidates, each had three programs meeting the exploratory prioritization criteria and stable negative donor-equal responses across tissues, whereas no source program met the global GSEA FDR threshold and no TF–program pair satisfied the full support rule. The corresponding predicted cross-tissue downregulated leading-edge genes therefore remain directional candidates. Applying the same framework to the myeloid TFs yielded no stable cross-tissue candidate. These results rank hypotheses for experimental perturbation without validating pathway inhibition, direct targets or causal mediation of a muscle–brain axis.

The stability analyses refined that prioritization. ETS2 and BHLHE40 retained concordant negative program responses across five mutually exclusive donor-group-out network rebuilds and additional parameter sensitivity analyses, while the state-space vector fields differed descriptively from randomized-residual controls. These checks test robustness within the fitted framework rather than biological replication. Donor sign tests quantify consistency of the custom program score; they do not replace the globally corrected GSEA results or demonstrate direct TF binding.

It should be noted that this study was designed to identify cross-tissue convergence and candidate regulatory responses rather than to establish causal muscle-to-brain signaling. In addition, several limitations should be considered. Muscle, brain and blood were drawn from independent public datasets, precluding direct assessment of within-person cross-organ coupling or mediation. The muscle comparison concerned age rather than clinically defined sarcopenia, and the small Frailty PBMC cohort represented a source-defined phenotype. Shared inflammatory pathways are not specific to a muscle–brain mechanism and may reflect common exposures or parallel responses. Differences in sequencing modality, cell recovery and statistical framework limit cross-dataset comparisons. Original counts were unavailable in the analyzed muscle object. The AD-spectrum PBMC analysis used the released ambient-RNA-corrected fractional-count matrix, whose processing must be considered when interpreting normalization and variance modeling. Age was unavailable in that PBMC analysis object, leaving potential residual confounding. The early CPS window was selected for cross-tissue comparison after tissue-specific characterization, and robustness to alternative boundaries was not established. Sparse endothelial representation in some brain donors and the small young-muscle myeloid sample further limit precision. CellChat used pooled cells or nuclei; bulk-blood expression was not resolved by cell type, and the frailty bulk analysis used RNA pools rather than individual libraries. CellOracle predictions remain conditional on the fitted networks and promoter prior, without matched chromatin-accessibility evidence. Conditional donor summaries do not capture all model uncertainty, and no candidate met the complete support rule. Replication in independent muscle and brain cohorts and experimental perturbation studies are needed.

## CONCLUSIONS

Independent human transcriptomic analyses identified shared endothelial and myeloid inflammatory programs in aging skeletal muscle and AD-related brain tissue. Related PBMC patterns and external pathway-level evidence from the frailty bulk-blood cohort and ADNI MCI/AD comparisons support the presence of related signals in peripheral blood. CellOracle simulations prioritized ETS2 and BHLHE40 as exploratory candidate regulators of the shared endothelial programs.

Together, these findings support a putative muscle–brain axis in AD characterized by convergent vascular and immune inflammatory programs. However, because the tissues were obtained from independent individuals and the analyses were cross-sectional, the results do not establish direct muscle-to-brain signaling, mediation, or causal coupling. Rather, they define specific cross-tissue programs and candidate regulators for testing in paired human cohorts and experimental models.

## LIST OF ABBREVIATIONS

AD: Alzheimer’s disease
ADNI: Alzheimer’s Disease Neuroimaging Initiative
APOE: apolipoprotein E
BH: Benjamini-Hochberg
CPM: counts per million
CPS: continuous pseudoprogression score
ECM: extracellular matrix
FAP: fibro/adipogenic progenitor
FDR: false discovery rate
GO BP: Gene Ontology Biological Process
GSEA: gene set enrichment analysis
HC: healthy control
HLMA: human limb skeletal muscle atlas
KO: knockout
L-R: ligand-receptor
MAST: Model-based Analysis of Single-cell Transcriptomics
MBA: muscle-brain axis
MCI: mild cognitive impairment
Micro-PVM: microglia/perivascular macrophage
MSigDB: Molecular Signatures Database
MTG: middle temporal gyrus
MuSC: muscle stem or satellite cell
NES: normalized enrichment score
NF-kappaB: nuclear factor kappa B
NK: natural killer
NKT: natural killer T
OPC: oligodendrocyte precursor cell
PBMC: peripheral blood mononuclear cell
scRNA-seq: single-cell RNA sequencing
SEA-AD: Seattle Alzheimer’s Disease Brain Cell Atlas
SMC: smooth muscle cell
snRNA-seq: single-nucleus RNA sequencing
TF: transcription factor
TMM: trimmed mean of M values
UMAP: uniform manifold approximation and projection
VLMC: vascular and leptomeningeal cell

## DECLARATIONS

## Ethics approval and consent to participate

This study was a secondary analysis of de-identified data obtained from publicly available or controlled-access repositories and involved no prospective recruitment, participant contact, intervention or new tissue collection. For the Seattle Alzheimer’s Disease Brain Cell Atlas (SEA-AD) resource, postmortem brain tissue and donor metadata were obtained by the source investigators through the University of Washington BioRepository and Integrated Neuropathology (BRaIN) Laboratory from participants in the Adult Changes in Thought study and the University of Washington Alzheimer’s Disease Research Center. The source study reports that informed consent for research brain donation was obtained under protocols approved by the University of Washington and Kaiser Permanente Washington Health Research Institute Institutional Review Boards, and that the work conducted at the Allen Institute received a determination of non-human-subject research. Ethical approval and informed consent for the other source cohorts were obtained by the original investigators as described in the respective source publications and repositories. Because the present work used only de-identified secondary data released for research use, ethics approval and consent to participate for the present analyses were not applicable.

## Consent for publication

Not applicable. This manuscript contains no identifiable individual participant information.

## Availability of data and materials

The datasets analyzed in this study were obtained from existing resources. The SEA-AD middle temporal gyrus analyses used the processed AnnData object SEAAD_MTG_RNAseq_final-nuclei.2026-06-22.h5ad (release dated 22 June 2026), available through the SEA-AD Data and Downloads portal (https://brain-map.org/consortia/sea-ad/our-data) and the Registry of Open Data on AWS (https://registry.opendata.aws/sea-ad-single-cell-profiling/). Raw SEA-AD single-nucleus sequencing data are available under controlled access through the AD Knowledge Portal (Synapse accession syn26223298). Access, use and redistribution of SEA-AD resources are subject to the Allen Institute Terms of Use and Citation Policy and any applicable external-partner terms. Processed skeletal-muscle transcriptomic data and metadata are publicly available through the Human skeletal muscle aging atlas (HLMA) portal (https://db.genomics.cn/cell/hlma/). Peripheral-blood datasets are publicly available from the Gene Expression Omnibus under accession numbers GSE157007 (https://www.ncbi.nlm.nih.gov/geo/query/acc.cgi?acc=GSE157007) and GSE206762 (https://www.ncbi.nlm.nih.gov/geo/query/acc.cgi?acc=GSE206762), and from the Sequence Read Archive under accession number SRP312418 (https://www.ncbi.nlm.nih.gov/sra/?term=SRP312418). The processed expression object and annotations associated with SRP312418 are available through the ADRCSC portal (https://www.ccb.uni-saarland.de/adrcsc). ADNI gene-expression and clinical data are accessible through the LONI Image and Data Archive (https://ida.loni.usc.edu/) following approval of a data-access application and acceptance of the ADNI Data Use Agreement, as described at https://adni.loni.usc.edu/data-samples/adni-data/. Individual-level ADNI data are not redistributed by the authors. All datasets remain subject to the access and reuse conditions of their respective providers. Supporting pathway-selection records, model summaries and analysis diagnostics are provided in Additional file 3 (Supplementary Tables S2–S6).

## Authors’ contributions

JZ (Jing Zhang) and WP conceived and designed the study. WS developed the methodology and performed the bioinformatics and statistical analyses. AZ curated the data and prepared the figures. AZ and WS drafted the manuscript. JC, XL, JFZ (Jingfeng Zou), SL, SH, CP, ZL, GP, QZ and FA interpreted the results and critically revised the manuscript. JZ, WP and SY (Shijun Yan) supervised the study. JZ acquired funding. All authors approved the manuscript for submission.

## Acknowledgements

We gratefully acknowledge the SEA-AD Consortium, the Allen Institute for Brain Science, the University of Washington BRaIN Laboratory, the Adult Changes in Thought study, the University of Washington Alzheimer’s Disease Research Center, and especially the study participants, brain donors and their families who made the SEA-AD resource possible. Generation of the SEA-AD resource was supported by the National Institute on Aging under awards U19AG060909, P50AG005136, P30AG066509, U01AG006781 and U19AG066567. Data used in the preparation of this article were obtained from the Alzheimer’s Disease Neuroimaging Initiative (ADNI) database (adni.loni.usc.edu). As such, investigators within ADNI contributed to the design and implementation of ADNI and/or provided data but did not participate in the analysis or writing of this report. A complete listing of ADNI investigators is available at http://adni.loni.usc.edu/wp-content/uploads/how_to_apply/ADNI_Acknowledgement_List.pdf. Data collection and sharing for ADNI are funded by the National Institute on Aging (National Institutes of Health Grant U19AG024904). The grantee organization is the Northern California Institute for Research and Education. In the past, ADNI has also received funding from the National Institute of Biomedical Imaging and Bioengineering, the Canadian Institutes of Health Research, and private-sector contributions through the Foundation for the National Institutes of Health, including contributions from AbbVie; Alzheimer’s Association; Alzheimer’s Drug Discovery Foundation; Araclon Biotech; BioClinica, Inc.; Biogen; Bristol-Myers Squibb Company; CereSpir, Inc.; Cogstate; Eisai Inc.; Elan Pharmaceuticals, Inc.; Eli Lilly and Company; EuroImmun; F. Hoffmann-La Roche Ltd and its affiliated company Genentech, Inc.; Fujirebio; GE Healthcare; IXICO Ltd.; Janssen Alzheimer Immunotherapy Research & Development, LLC; Johnson & Johnson Pharmaceutical Research & Development LLC; Lumosity; Lundbeck; Merck & Co., Inc.; Meso Scale Diagnostics, LLC; NeuroRx Research; Neurotrack Technologies; Novartis Pharmaceuticals Corporation; Pfizer Inc.; Piramal Imaging; Servier; Takeda Pharmaceutical Company; and Transition Therapeutics.

We also acknowledge the investigators, study participants and data stewards who made the HLMA, GSE157007, GSE206762, SRP312418 and ADRCSC resources available.

## REFERENCES

1. Franceschi C, Garagnani P, Parini P, Giuliani C, Santoro A. Inflammaging: a new immune–metabolic viewpoint for age-related diseases. Nat Rev Endocrinol. 2018;14:576–90. 10.1038/s41574-018-0059-4

2. Boyle PA, Buchman AS, Wilson RS, Leurgans SE, Bennett DA. Association of muscle strength with the risk of alzheimer disease and the rate of cognitive decline in community-dwelling older persons. Arch Neurol. 2009;66:1339–44. 10.1001/archneurol.2009.240

3. McNeish BL, Miljkovic I, Liu-Ambrose T, Ambrosio F, Esser K, Fahnestock M, et al. Muscle-brain crosstalk as a driver of brain health in aging. GeroScience. 2025;48:4873–91. 10.1007/s11357-025-01833-0

4. Lai Y, Ramírez-Pardo I, Isern J, An J, Perdiguero E, Serrano AL, et al. Multimodal cell atlas of the ageing human skeletal muscle. Nature. 2024;629:154–64. 10.1038/s41586-024-07348-6

5. Kedlian VR, Wang Y, Liu T, Chen X, Bolt L, Tudor C, et al. Human skeletal muscle aging atlas. Nat Aging. 2024;4:727–44. 10.1038/s43587-024-00613-3

6. Mathys H, Davila-Velderrain J, Peng Z, Gao F, Mohammadi S, Young JZ, et al. Single-cell transcriptomic analysis of Alzheimer’s disease. Nature. 2019;570:332–7. 10.1038/s41586-019-1195-2

7. Yang AC, Vest RT, Kern F, Lee DP, Agam M, Maat CA, et al. A human brain vascular atlas reveals diverse mediators of Alzheimer’s risk. Nature. 2022;603:885–92. 10.1038/s41586-021-04369-3

8. Sun N, Victor MB, Park YP, Xiong X, Scannail AN, Leary N, et al. Human microglial state dynamics in Alzheimer’s disease progression. Cell. 2023;186:4386–4403.e29. 10.1016/j.cell.2023.08.037

9. Amersfoort J, Eelen G, Carmeliet P. Immunomodulation by endothelial cells — partnering up with the immune system? Nat Rev Immunol. 2022;22:576–88. 10.1038/s41577-022-00694-4

10. Grandke F, Fehlmann T, Kern F, Gate DM, Wolff TW, Leventhal O, et al. A single-cell atlas to map sex-specific gene-expression changes in blood upon neurodegeneration. Nat Commun. 2025;16:1965. 10.1038/s41467-025-56833-7

11. Kamimoto K, Stringa B, Hoffmann CM, Jindal K, Solnica-Krezel L, Morris SA. Dissecting cell identity via network inference and in silico gene perturbation. Nature. 2023;614:742–51. 10.1038/s41586-022-05688-9

12. Hao Y, Stuart T, Kowalski MH, Choudhary S, Hoffman P, Hartman A, et al. Dictionary learning for integrative, multimodal and scalable single-cell analysis. Nat Biotechnol. 2024;42:293–304. 10.1038/s41587-023-01767-y

13. Benjamini Y, Hochberg Y. Controlling the False Discovery Rate: A Practical and Powerful Approach to Multiple Testing. Journal of the Royal Statistical Society Series B: Statistical Methodology. 1995;57:289–300. 10.1111/j.2517-6161.1995.tb02031.x

14. Squair JW, Gautier M, Kathe C, Anderson MA, James ND, Hutson TH, et al. Confronting false discoveries in single-cell differential expression. Nat Commun. 2021;12:5692. 10.1038/s41467-021-25960-2

15. Zimmerman KD, Espeland MA, Langefeld CD. A practical solution to pseudoreplication bias in single-cell studies. Nat Commun. 2021;12:738. 10.1038/s41467-021-21038-1

16. Robinson MD, McCarthy DJ, Smyth GK. edgeR : a Bioconductor package for differential expression analysis of digital gene expression data. Bioinformatics. 2010;26:139–40. 10.1093/bioinformatics/btp616

17. Robinson MD, Oshlack A. A scaling normalization method for differential expression analysis of RNA-seq data. Genome Biol. 2010;11:R25. 10.1186/gb-2010-11-3-r25

18. Ritchie ME, Phipson B, Wu D, Hu Y, Law CW, Shi W, et al. limma powers differential expression analyses for RNA-sequencing and microarray studies. Nucleic Acids Research. 2015;43:e47–e47. 10.1093/nar/gkv007

19. Subramanian A, Tamayo P, Mootha VK, Mukherjee S, Ebert BL, Gillette MA, et al. Gene set enrichment analysis: A knowledge-based approach for interpreting genome-wide expression profiles. Proc Natl Acad Sci USA. 2005;102:15545–50. 10.1073/pnas.0506580102

20. Wu T, Hu E, Xu S, Chen M, Guo P, Dai Z, et al. clusterProfiler 4.0: A universal enrichment tool for interpreting omics data. The Innovation. 2021;2:100141. 10.1016/j.xinn.2021.100141

21. Liberzon A, Birger C, Thorvaldsdóttir H, Ghandi M, Mesirov JP, Tamayo P. The Molecular Signatures Database Hallmark Gene Set Collection. Cell Systems. 2015;1:417–25. 10.1016/j.cels.2015.12.004

22. Kanehisa M, Furumichi M, Sato Y, Matsuura Y, Ishiguro-Watanabe M. KEGG: biological systems database as a model of the real world. Nucleic Acids Research. 2025;53:D672–7. 10.1093/nar/gkae909

23. The Gene Ontology Consortium, Aleksander SA, Balhoff JP, Carbon S, Cherry JM, Ebert D, et al. The Gene Ontology knowledgebase in 2026. Nucleic Acids Research. 2026;54:D1779–92. 10.1093/nar/gkaf1292

24. Kolmykov S, Yevshin I, Kulyashov M, Sharipov R, Kondrakhin Y, Makeev VJ, et al. GTRD: an integrated view of transcription regulation. Nucleic Acids Research. 2021;49:D104–11. 10.1093/nar/gkaa1057

25. Wang JZ, Du Z, Payattakool R, Yu PS, Chen C-F. A new method to measure the semantic similarity of GO terms. Bioinformatics. 2007;23:1274–81. 10.1093/bioinformatics/btm087

26. Jin S, Guerrero-Juarez CF, Zhang L, Chang I, Ramos R, Kuan C-H, et al. Inference and analysis of cell-cell communication using CellChat. Nat Commun. 2021;12:1088. 10.1038/s41467-021-21246-9

27. Jin S, Plikus MV, Nie Q. CellChat for systematic analysis of cell–cell communication from single-cell transcriptomics. Nat Protoc. 2025;20:180–219. 10.1038/s41596-024-01045-4

28. Gabitto MI, Travaglini KJ, Rachleff VM, Kaplan ES, Long B, Ariza J, et al. Integrated multimodal cell atlas of Alzheimer’s disease. Nat Neurosci. 2024;27:2366–83. 10.1038/s41593-024-01774-5

29. He L, Davila-Velderrain J, Sumida TS, Hafler DA, Kellis M, Kulminski AM. NEBULA is a fast negative binomial mixed model for differential or co-expression analysis of large-scale multi-subject single-cell data. Commun Biol. 2021;4:629. 10.1038/s42003-021-02146-6

30. Milacic M, Beavers D, Conley P, Gong C, Gillespie M, Griss J, et al. The Reactome Pathway Knowledgebase 2024. Nucleic Acids Research. 2024;52:D672–8. 10.1093/nar/gkad1025

31. Lambert SA, Jolma A, Campitelli LF, Das PK, Yin Y, Albu M, et al. The human transcription factors. Cell. 2018;172:650–65. 10.1016/j.cell.2018.01.029

32. Finak G, McDavid A, Yajima M, Deng J, Gersuk V, Shalek AK, et al. MAST: a flexible statistical framework for assessing transcriptional changes and characterizing heterogeneity in single-cell RNA sequencing data. Genome Biol. 2015;16:278. 10.1186/s13059-015-0844-5

33. Wu D, Smyth GK. Camera: a competitive gene set test accounting for inter-gene correlation. Nucleic Acids Res. 2012;40:e133–e133. 10.1093/nar/gks461

34. Luo OJ, Lei W, Zhu G, Ren Z, Xu Y, Xiao C, et al. Multidimensional single-cell analysis of human peripheral blood reveals characteristic features of the immune system landscape in aging and frailty. Nat Aging. 2022;2:348–64. 10.1038/s43587-022-00198-9

35. Korsunsky I, Millard N, Fan J, Slowikowski K, Zhang F, Wei K, et al. Fast, sensitive and accurate integration of single-cell data with Harmony. Nat Methods. 2019;16:1289–96. 10.1038/s41592-019-0619-0

36. Büttner M, Ostner J, Müller CL, Theis FJ, Schubert B. scCODA is a Bayesian model for compositional single-cell data analysis. Nat Commun. 2021;12:6876. 10.1038/s41467-021-27150-6

37. Law CW, Chen Y, Shi W, Smyth GK. voom: precision weights unlock linear model analysis tools for RNA-seq read counts. Genome Biol. 2014;15:R29. 10.1186/gb-2014-15-2-r29

38. Chen Y, Lun ATL, Smyth GK. From reads to genes to pathways: differential expression analysis of RNA-seq experiments using rsubread and the edgeR quasi-likelihood pipeline. F1000Research. 2016;5:1438. 10.12688/f1000research.8987.2

39. Iparraguirre L, Alberro A, Iñiguez SG, Muñoz-Culla M, Vergara I, Matheu A, et al. Blood RNA-Seq profiling reveals a set of circular RNAs differentially expressed in frail individuals. Immun Ageing. 2023;20:33. 10.1186/s12979-023-00356-6

40. Agrawal A, Balcı H, Hanspers K, Coort SL, Martens M, Slenter DN, et al. WikiPathways 2024: next generation pathway database. Nucleic Acids Research. 2024;52:D679–89. 10.1093/nar/gkad960

41. Saykin AJ, Shen L, Yao X, Kim S, Nho K, Risacher SL, et al. Genetic studies of quantitative MCI and AD phenotypes in ADNI: Progress, opportunities, and plans. Alzheimer’s & Dementia. 2015;11:792–814. 10.1016/j.jalz.2015.05.009

42. Lukjanenko L, Karaz S, Stuelsatz P, Gurriaran-Rodriguez U, Michaud J, Dammone G, et al. Aging Disrupts Muscle Stem Cell Function by Impairing Matricellular WISP1 Secretion from Fibro-Adipogenic Progenitors. Cell Stem Cell. 2019;24:433–446.e7. 10.1016/j.stem.2018.12.014

43. Hayden MS, Ghosh S. NF-κB in immunobiology. Cell Res. 2011;21:223–44. 10.1038/cr.2011.13

44. Villarino AV, Kanno Y, O’Shea JJ. Mechanisms and consequences of Jak–STAT signaling in the immune system. Nat Immunol. 2017;18:374–84. 10.1038/ni.3691

45. Gate D, Saligrama N, Leventhal O, Yang AC, Unger MS, Middeldorp J, et al. Clonally expanded CD8 T cells patrol the cerebrospinal fluid in Alzheimer’s disease. Nature. 2020;577:399–404. 10.1038/s41586-019-1895-7

46. Cruz-Jentoft AJ, Bahat G, Bauer J, Boirie Y, Bruyère O, Cederholm T, et al. Sarcopenia: revised European consensus on definition and diagnosis. Age and Ageing. 2019;48:16–31. 10.1093/ageing/afy169

