## Supplemental_File_1_Supplementary_Methods for "Cross-Tissue Inflammatory Convergence Supports a Putative Muscle–Brain Axis in Alzheimer’s Disease"

Contents

SUPPLEMENTARY METHODS SCOPE

FIGURE DISPLAY CONVENTIONS

MUSCLE CELLCHAT IMPLEMENTATION

SEA-AD PATHWAY DISPLAY CURATION

SEA-AD IMMUNE COMPARTMENT DISPLAY

DONOR HEATMAP CONSTRUCTION

MUSCLE MYELOID MAST-GSEA SENSITIVITY DISPLAY

PBMC VISUALIZATION AND ANNOTATION HARMONIZATION

PBMC COMPOSITIONAL AND PSEUDOBULK IMPLEMENTATION

PBMC PATHWAY AND GENE DISPLAY SELECTION

BULK-BLOOD CROSS-MODALITY ANALYSES

CELLORACLE CANDIDATES AND ANALYSIS POPULATIONS

STATE DEFINITIONS AND CLUSTER-SPECIFIC REGULATORY MODELS

PROGRAM RESPONSES AND MULTIPLE-TESTING FAMILIES

CROSS-TISSUE EVIDENCE AUDIT AND SENSITIVITY ANALYSES

STATE-SPACE VECTORS AND RANDOMIZED-RESIDUAL CONTROLS

FIGURE 6 CORE PROGRAM AND LEADING-EDGE GENE SELECTION

SUPPLEMENTARY FIGURES S9 AND S10 ORGANIZATION

INTERPRETATION BOUNDARIES

Supplementary Table S1. Characteristics of the analyzed muscle and brain donors

**SUPPLEMENTARY METHODS SCOPE**

These procedures provide implementation and display details for Figures 1-6 and Supplementary Figures S1-S10. Cohort definitions, biological replicate units, statistical models, multiple-testing families and interpretation boundaries follow the main Methods. Reference numbers refer to the main manuscript bibliography. Display-specific filtering did not replace the complete source testing families or alter the reported effect estimates, P values or false discovery rates. Additional file 3 contains the pathway-selection records (Table S2), SEA-AD model summaries and GSEA source tables (Table S3), MAST diagnostics and sensitivity results (Table S4), CellOracle evidence and ranking tables (Table S5), and PBMC matrix-scale verification (Table S6).

**FIGURE DISPLAY CONVENTIONS**

Figures used Arial and a white background. Age groups used colorblind-safe blue for Young (#0072B2) and orange for Old (#D55E00); muscle perturbation annotations used burgundy and brain annotations used blue-gray. Gene symbols were italicized where applicable, whereas pathway and cell-type names were set in roman type. Blank heatmap positions indicate unavailable combinations, and gray perturbation cells or placeholders indicate that a virtual knockout was not estimable from the fitted network rather than evidence of no biological effect.

**MUSCLE CELLCHAT IMPLEMENTATION**

Separate CellChat objects were constructed from all cells pooled within the Young (86,588 cells) and Old (205,835 cells) groups [38,39]. Analyses used CellChatDB.human, trimean average expression, computeCommunProb with 100 bootstrap iterations, and filterCommunication with min.cells = 10, followed by computeCommunProbPathway, aggregateNet and netAnalysis_computeCentrality. The objects were combined with mergeCellChat. Differential communication strength was defined as the Old-minus-Young difference in the weighted networks (@net$weight). Figure 1f displays this sender-receiver matrix. Pathway comparisons used rankNet with mode = "comparison", measure = "weight", stacked = TRUE and do.stat = TRUE. These group-pooled network comparisons were treated as model-based descriptive inference rather than donor-level tests.

**SEA-AD PATHWAY DISPLAY CURATION**

SEA-AD analyses used the processed middle temporal gyrus single-nucleus RNA-sequencing AnnData object SEAAD_MTG_RNAseq_final-nuclei.2026-06-22.h5ad (release dated 22 June 2026), obtained from the SEA-AD Data and Downloads portal and the Registry of Open Data on AWS [11]. Donor metadata, source cell annotations, two-dimensional embeddings and the continuous pseudoprogression score (CPS) were taken from the same release; raw sequence reads were not reprocessed. The released object contained 1,178,694 nuclei from 84 donors aged 65–102 years. The full cohort was used for descriptive displays, whereas inferential analyses used 73 donors after exclusion of the 11 source-designated severely affected donors. Early- and late-CPS-window definitions and their overlap are reported in the main Methods and Supplementary Table S1.

The early- and late-CPS-window displays used pathways from the Hallmark, KEGG Legacy and Gene Ontology Biological Process collections. GO Biological Process terms were greedily reduced at a Wang semantic-similarity threshold of 0.7 [37], after which related terms were organized into prespecified biological themes. Figure 2f retains 16 early-window pathways carried forward for the main comparison: 10 inflammation or immunity pathways and six extracellular-matrix, adhesion or fibrosis pathways. Supplementary Figure S3 displays 34 late-window pathways across inflammation or immunity; cytoskeleton, extracellular matrix or vascular biology; signaling or stress response; cell growth or proliferation; and metabolism. Every displayed pathway reached FDR < 0.05 in at least one analysis unit within the corresponding window. Display curation did not alter the source normalized enrichment scores or FDR values.

Supplementary Table S2a preserves the original 33-pathway muscle curation pool, its biological themes and membership in the 16-pathway Figure 1e display. The other 17 pathways remained in the original results and were omitted only from the focused display. Table S2b preserves the 112 pathway-by-cell-type records in the final 16-pathway early SEA-AD display; Tables S2c and S2d contain the late-window curation and saved GO redundancy decisions, respectively. These display records do not define the multiple-testing families.

For the seven principal SEA-AD analysis units, the archived model outputs provide actual donor and nucleus counts, gene-level statistics and convergence codes (Table S3a). Lymphocyte models used 28 donors and 150 nuclei in the early window and 45 donors and 534 nuclei in the late window; the other six units retained all 30 early-window and 48 late-window donors. Missing P values or effect statistics were retained as unavailable, without imputation. Gene FDR values corresponded to Benjamini-Hochberg adjustment of the available P values within each unit-by-window model. The four archived GSEA collection exports retain their original normalized enrichment scores and adjusted P values (Table S3b).

In the immune and endothelial GSEA workflows, genes with missing P values or nonfinite z statistics were excluded from ranking; duplicate symbols were represented by the z statistic with the largest absolute value. Enrichment tasks required at least 1,000 rankable genes. fgseaMultilevel used gene sets with 10-500 represented genes, gseaParam = 1, eps = 0 and seed 42. Hallmark, GO BP, Reactome and KEGG Legacy were combined in one enrichment-testing family for each unit and window, whereas TFT GTRD was analyzed separately. Source adjusted P values were retained during display selection.

**SEA-AD IMMUNE COMPARTMENT DISPLAY**

Supplementary Figure S2 defines the immune populations used in the brain analyses. Twelve Monocyte nuclei were excluded because this source label was too sparse for downstream modeling. The retained immune compartment contained 36,674 nuclei assigned to six Microglia/perivascular macrophage (Micro-PVM) supertypes and Lymphocyte. The local immune UMAP retained the source embedding without trajectory refitting. Twelve canonical identity markers were cross-checked against data-derived marker rankings and displayed across the seven retained supertypes. Downstream Micro-PVM combined the six myeloid supertypes and excluded Lymphocyte and Monocyte. The display supported annotation and analysis scope only and did not introduce a new between-group test.

**DONOR HEATMAP CONSTRUCTION**

Brain counts were summed by donor and analysis unit and converted to log1p counts per million. Endothelial heatmaps included 693 nuclei from 30 early-window donors, and Micro-PVM heatmaps included 13,416 nuclei from the same 30 donors. Muscle endothelial heatmaps included 23,227 cells from 23 donors (8 Young and 15 Old), whereas the muscle myeloid heatmaps included 259 nuclei from 4 Young donors and 4,166 nuclei from 9 Old donors. Columns were ordered by age in muscle and CPS in brain. Expression was standardized separately by gene within each tissue and displayed from -2 to 2 without row or column clustering. Five brain donors with fewer than 10 endothelial nuclei were marked as a display-quality annotation rather than excluded.

The Figure 3 muscle side bars show donor-aggregate limma log2 fold changes, and the brain side bars show NEBULA log2 fold changes per 0.1 CPS. In Figure 4, muscle side bars show the MAST-adjusted predicted mean log2-expression difference and brain side bars show the corresponding NEBULA effect per 0.1 CPS. Effect-bar lengths were not compared across tissues. The complete 109-gene endothelial and 51-gene myeloid heatmaps are shown in Supplementary Figures S4 and S5, respectively. Leading-edge membership was not interpreted as gene-level significance in both tissues.

**MUSCLE MYELOID MAST-GSEA SENSITIVITY DISPLAY**

The muscle myeloid MAST analysis was restricted to Chinese single-nucleus donors with at least 20 myeloid nuclei and retained 4 Young and 9 Old donors. Supplementary Figure S1 displays 9 of the 16 Figure 1e reference pathways that reached nominal MAST-GSEA P < 0.05: five inflammation or immunity pathways and four extracellular-matrix, adhesion or fibrosis pathways. Dark outlines indicate the original complete-family FDR < 0.05. Focal adhesion and tight junction met the nominal display threshold but not the database-wide FDR threshold. This supplementary display rule was not used to select the shared pathways in Figure 4.

The fixed testing set comprised 6,383 genes detected in at least 10% of myeloid nuclei in at least three included donors and with variable donor-mean expression. The source log1p-normalized matrix was divided by log(2) to obtain log2(1 + normalized expression). Cellular detection rate used all 48,355 source features and was standardized across included nuclei without age-group labels. MAST 1.38.0 with the lme4 2.0-6 mixed-model backend fitted ~ age_group + Sex + cdr_z + (1 | donor), with Young as the reference. Both hurdle components included a donor random intercept. The continuous component used maximum likelihood (REML = FALSE), empirical Bayes shrinkage was disabled, and the discrete component used the Laplace approximation (nAGQ = 1). Fits first used bobyqa; invalid fits were retried with a predefined nloptwrap configuration rather than selected by statistical significance.

The full model was compared with the model omitting age group using a joint likelihood-ratio test. Validity required convergence of each required full and reduced component, a zero optimizer error code, no convergence messages or dropped coefficients, finite effect and joint-test statistics, and nonnegative component likelihood-ratio statistics within numerical tolerance. Random-effect boundaries were flagged with lme4::isSingular(tol = 1e-4) in any full or reduced component and were recorded separately from numerical failure. Of 6,383 planned gene tests, 6,362 had valid fits, including 4,389 boundary fits; 21 unresolved fits were excluded from inference and ranking. Gene-level Benjamini-Hochberg correction retained all 6,383 planned tests as the family size. Component-specific effects and diagnostics are provided in Table S4a.

GSEA used the signed square root of the joint statistic, oriented by the donor-equally weighted adjusted Old-minus-Young mean difference; this ranking is not a standard-normal z statistic. The Hallmark, KEGG Legacy and GO BP collections were tested separately with fgseaMultilevel (minSize = 10, maxSize = 500, eps = 1e-8, nPermSimple = 5,000, sampleSize = 101; seed 20260919). Numerically unresolved pathway estimates were retried with nPermSimple = 100,000 and seed 20261919, followed by Benjamini-Hochberg correction over the complete eligible collection. MAST and donor-level limma were compared on the same 6,362-gene valid-fit universe and, separately, on the 1,973-gene universe excluding boundary fits. All four Figure 4 inflammatory pathways retained positive MAST enrichment scores after boundary exclusion, but only inflammatory response retained FDR < 0.05. The reduced universe also changes gene-set coverage, so differences cannot be attributed solely to boundary status. Complete results, including CAMERA and donor pathway-score tests, are provided in Table S4. These analyses reuse the same donors and constitute sensitivity analyses.

**PBMC VISUALIZATION AND ANNOTATION HARMONIZATION**

Figure 5a displays all 76,509 quality-controlled GSE157007 peripheral blood mononuclear cells from six healthy older controls and five participants with the source-defined Frailty phenotype. The source Harmony embedding was retained (30 Harmony dimensions, 30 neighbors, minimum distance 0.30, cosine metric and seed 42), and diagnosis or age was not used to generate the embedding [42]. Figure 5b displays a deterministic diagnosis- and cell-type-stratified subset of 250,096 cells from the 696,093-cell SRP312418 HC/MCI/AD atlas. Published SRP312418 cell-type and diagnosis endpoints lacked shared barcodes but contained unique matching UMAP-coordinate pairs; these pairs were joined one-to-one for visualization only. Parkinson disease and Parkinson disease with MCI cells were excluded. The two atlases were not jointly embedded, and display sampling did not affect composition or pseudobulk analyses.

Source labels were harmonized only at a broad biological level. The six main PBMC analysis populations were monocytes, CD4+ T cells, CD8+ T cells, B cells, NK cells and NKT-like cells. NKT-like/T-NK* aligns broad source labels and does not establish identical subpopulations. Frailty denotes the phenotype defined by the source study and was not reclassified as clinically diagnosed sarcopenia.

**PBMC COMPOSITIONAL AND PSEUDOBULK IMPLEMENTATION**

Cell-type proportions used all quality-controlled PBMCs from each participant as the denominator. Bootstrap percentile 95% confidence intervals for unadjusted mean percentage-point differences used 2,000 participant resamples for SRP312418 and 10,000 for GSE157007. Primary compositional inference used scCODA [43]. The SRP312418 model included diagnosis and sex, used plasma cells as the reference, and used 1,000 warm-up and 2,000 posterior samples at an estimated FDR of 5%. The GSE157007 model included group, standardized age and source batch, used CD4+ T cells as the reference, and used 1,000 warm-up and 5,000 posterior samples at an estimated FDR of 5%. Bootstrap intervals were descriptive and did not replace the scCODA posterior-selection rule.

SRP312418 participant-by-cell-type pseudobulks required at least 20 cells and at least 10 retained participants per group. The released H5AD X matrix contained non-negative fractional counts on the linear expression scale. Differential-expression analysis used TMM normalization, voom precision weights and robust limma empirical Bayes moderation with diagnosis and sex [28,29,44]. GSE157007 used integer-verified RNA/counts data, at least 10 cells per pseudobulk and at least three participants per group, followed by TMM normalization and edgeR robust quasi-likelihood modeling with source batch, group and standardized age [27]. Participants, rather than cells, were the biological replicates.

The SRP312418 source study applied SoupX 1.4.8 to correct ambient RNA before downstream expression analysis. In the released ADRC_AnnotatedObject.h5ad, X was a 909,322-cell by 55,932-gene sparse matrix with no additional raw-count layer. A complete read-only check found no negative or nonfinite entries. X row sums reproduced nCount_RNA for every cell (maximum absolute difference, 1.10 x 10^-10), and positive-feature counts reproduced nFeature_RNA for all 909,322 cells, consistent with the source study's corrected-count representation (Table S6). The analysis summed these fractional values by participant and cell type without rounding, exponentiation or additional per-cell normalization. Design-aware gene filtering used edgeR::filterByExpr (min.count = 10, min.total.count = 15) before TMM normalization and limma-voom. For GSE157007, filterByExpr used the design matrix and min.count = 10. Gene-level Benjamini-Hochberg correction was applied separately within each cell-type-by-contrast analysis.

**PBMC PATHWAY AND GENE DISPLAY SELECTION**

PBMC GSEA used the frozen MSigDB 2026.1.Hs Hallmark collection and fgseaMultilevel, retaining gene sets represented by 15-500 ranked genes. SRP312418 genes were ranked by the moderated t statistic and GSE157007 genes by sign(log2 fold change) multiplied by the square root of the quasi-likelihood F statistic. Antibody-derived features were excluded. Within each contrast, Benjamini-Hochberg correction was applied across all eligible Hallmark pathway-by-cell-type combinations in the six main PBMC populations as one prespecified family. Figure 5d displays the fixed six source-related Hallmark pathways without recalculating FDR over that subset.

Figures 5e and 5f project the complete 109-gene endothelial and 51-gene myeloid tissue intersections into PBMC differential-expression results. A gene was retained for display when log2 fold change was positive and unadjusted P < 0.05 in at least one displayed cell-type-by-contrast context, yielding 61 endothelial-source and 32 myeloid-source genes. All available tests for retained genes were represented using the stated display encoding; pale cells do not show the magnitude of effects that failed the positive nominal rule. Gene-level FDR values came from the complete original testing family and were not recalculated over the displayed genes. Crosses denote unavailable tests, whereas pale cells denote available results that did not meet the positive nominal display rule. Endothelial-source and myeloid-source identify the tissue intersection from which a gene was selected and do not assign a circulating cell of origin.

**BULK-BLOOD CROSS-MODALITY ANALYSES**

Data used for the ADNI analyses were obtained from the Alzheimer's Disease Neuroimaging Initiative (ADNI) database (adni.loni.usc.edu). ADNI was launched in 2003 as a public-private partnership led by Principal Investigator Michael W. Weiner, MD. Its original goal was to test whether serial magnetic resonance imaging, positron emission tomography, other biological markers, and clinical and neuropsychological assessments could be combined to measure the progression of mild cognitive impairment and early Alzheimer's disease. Current goals include validating biomarkers for clinical trials, improving the generalizability of ADNI data by increasing cohort diversity, and providing data on the diagnosis and progression of Alzheimer's disease to the scientific community. The present study used PAXgene whole-blood Affymetrix microarray expression data and linked clinical and demographic variables from the ADNI GO and ADNI2 phases; individual-level ADNI data were accessed under the ADNI Data Use Agreement and were not redistributed by the authors. For up-to-date information, see adni.loni.usc.edu.

GSE206762 analyzed red-blood-cell-lysed peripheral-blood leukocyte RNA sequencing from 20 sex-stratified RNA pools, with 10 pools per Frail or Robust phenotype; pools, rather than the 70 contributing participants, were the statistical replicates [13]. ADNI whole-blood microarray analyses included 244 HC, 376 MCI and 112 AD participants, with MCI and AD each compared with the common HC group after adjustment for age, sex, RNA integrity number, study phase and array plate [14]. Effects were not pooled across RNA-sequencing and microarray platforms.

Supplementary Figure S6 tested pathway-specific subsets fixed from the Figure 3 endothelial and Figure 4 myeloid intersections against each complete bulk ranking. Benjamini-Hochberg correction covered the prespecified 30 source-by-pathway-by-comparison tests. Complement and TNF-alpha/NF-kappaB were undefined for the myeloid source because Figure 4 supplied no corresponding intersection subset. Supplementary Figures S7 and S8 display all available effects for the 38 of 109 endothelial-source and 18 of 51 myeloid-source genes that were nominally increased in at least one bulk comparison. None reached gene-level FDR < 0.05 in any bulk dataset. Open circles therefore indicate nominal P < 0.05 and were interpreted as exploratory directional agreement.

**CELLORACLE CANDIDATES AND ANALYSIS POPULATIONS**

The Figure 3 intersection supplied 18 endothelial transcription-factor candidates, and the Figure 4 intersection supplied six myeloid candidates: IRF1, STAT3, SPI1, STAT2, MXD1 and PLSCR1. Endothelial models used 2,753 HLMA profiles from 15 Old donors and 640 SEA-AD endothelial nuclei from 30 early-CPS-window donors. Myeloid models used 471 HLMA single-nucleus profiles from eight Old Chinese donors and 8,200 SEA-AD Micro-PVM nuclei from the same 30 early-window donors. Each donor contributed at most 300 randomly sampled profiles (seed 20260921). Eligible states contained at least 100 profiles and at least five donors contributing three or more profiles each.

A tissue-level virtual knockout was estimable when the candidate passed expression and promoter-prior screening and at least one eligible state contained at least five fitted outgoing TF-target edges. One to four edges were retained as an insufficient-support category, and zero edges were recorded separately. Fifteen endothelial candidates were estimable in at least one tissue and 13 in both; PLSCR1, RBCK1 and TRAFD1 were not estimable in either tissue. IRF1, STAT3, SPI1 and STAT2 were estimable in both myeloid tissues, whereas MXD1 and PLSCR1 were not. Non-estimability was not interpreted as evidence of no biological function.

**STATE DEFINITIONS AND CLUSTER-SPECIFIC REGULATORY MODELS**

Muscle endothelial states were described as Capillary-like, Stress response, Venous-like and Remodeling-like from marker evidence. These labels were descriptive or provisional, and the Stress response state was strongly enriched for scRNA profiles. Brain Endo_2 and Endo_3 were displayed as BBB capillary-like and Inflammatory venular-like while retaining the source state identities. Muscle myeloid states were described provisionally as Activation-like and Glycolytic inflammatory-like; brain myeloid analyses retained five Microglia/PVM source states. These names aided biological interpretation and did not define a new taxonomy, pseudotime or developmental trajectory.

For each lineage, 3,000 shared expression-supported features were fixed while retaining eligible candidate TFs. CellOracle v0.20.0 used the public hg38 promoter Base-GRN without matched cell-type-specific chromatin-accessibility data [10]. State-specific networks were fitted by ridge regression with alpha = 10 and 20 rounds of cell bagging. Edges were filtered at model P <= 0.001 and by absolute coefficient, retaining at most 2,000 edges per state. Each estimable TF was set to zero and its predicted response propagated for three steps. The resulting changes were predictions within the fitted feature space and were not measurements from experimental knockout cells.

Shared features were required to be detected in at least 10 profiles and 1% of profiles in each tissue and were ranked by the mean of their tissue-specific normalized-dispersion percentile ranks. Before network fitting, CellOracle applied balanced k-nearest-neighbor imputation to normalized log1p expression over 30 principal components, with k equal to 2.5% of profiles, rounded to the nearest integer, and a minimum of 10. The pre-fit TF screen required expression in at least 10 profiles and 1% of profiles, at least three donors with three or more expressing profiles each, and at least five promoter-prior targets among the selected features.

**PROGRAM RESPONSES AND MULTIPLE-TESTING FAMILIES**

Predicted knockout-minus-wild-type changes were summarized within donors and averaged with equal donor weights. Source programs comprised the six Figure 3 endothelial inflammatory programs or four Figure 4 myeloid programs. Scores excluded the directly zeroed TF and averaged member-gene delta log1p values divided by the baseline standard deviation, with a standard-deviation floor of 0.1 and at least 10 modeled members required. Conditional 95% intervals used 2,000 donor bootstrap resamples. Exact two-sided donor sign tests were Benjamini-Hochberg-corrected across the corresponding source programs within each knockout. These q values summarize directional consistency of the custom score conditional on shared fitted networks and are distinct from global GSEA FDR. They do not capture uncertainty from network estimation.

Perturbation GSEA ranked donor-equal predicted delta log1p values after excluding the perturbed TF and used the frozen Hallmark 2026.1.Hs collection. Global Benjamini-Hochberg correction was applied within each tissue and lineage across all estimable TFs and eligible Hallmark tests. This global family, rather than the within-knockout sign-test family, determined whether a source pathway showed multiplicity-corrected predicted suppression.

**CROSS-TISSUE EVIDENCE AUDIT AND SENSITIVITY ANALYSES**

The cross-tissue evidence audit recorded program-score and conditional-interval signs, GSEA direction and global FDR, five mutually exclusive donor-group-out network rebuilds, and six edge-filter or propagation variants. The endothelial audit also required concordance in a muscle snRNA-only reconstruction. A fully supported TF-program pair required negative responses and intervals in both tissues, global GSEA FDR < 0.05 in both, and concordance across all applicable robustness checks. Rebuilds repeated imputation, bagged network inference, edge filtering and propagation with fixed features and state labels. They used their own baseline standard deviations and were treated as sensitivity analyses rather than independent biological replicates.

An exploratory-priority TF-program pair required a negative donor-equal score and an upper conditional 95% confidence limit below zero in both tissues, concordant negative responses in all five donor-group-out rebuilds and all six parameter variants in each tissue, and negative GSEA normalized enrichment scores in both tissues. Endothelial pairs additionally required a negative score in the muscle snRNA-only reconstruction. The corresponding stable-negative category required all of these conditions except the GSEA-direction criterion. Global GSEA FDR < 0.05 in both tissues was an additional requirement for the stricter fully supported category, not for exploratory prioritization.

Endothelial candidates were ordered by decreasing number of exploratory-priority programs, then by the numbers of stable-negative programs, programs with negative scores in both tissues, and programs with negative GSEA scores in both tissues. Remaining ties used the largest, across source programs, of the smaller absolute muscle and brain program scores. ETS2 and BHLHE40 each had three exploratory-priority programs and ranked first and second, respectively. No endothelial or myeloid pair met the fully supported rule. Table S5 provides the complete evidence fields and endothelial ordering; these descriptive priorities do not establish multiplicity-corrected pathway inhibition.

**STATE-SPACE VECTORS AND RANDOMIZED-RESIDUAL CONTROLS**

For each TF estimable in at least one tissue, CellOracle transition probabilities were projected onto the observed two-dimensional state space. A native randomized-residual calculation was processed through the same projection pipeline as a descriptive control. Within each tissue, virtual-knockout and randomized maps used identical grid points, thinning, arrow scale, color, width and opacity. Scales were not compared across the independent muscle and brain embeddings. Grid points are display positions rather than biological replicates, and a single randomized field does not provide a biological P value or null distribution. No pseudotime root or Young-state trajectory was imposed.

**FIGURE 6 CORE PROGRAM AND LEADING-EDGE GENE SELECTION**

Figure 6e retained three selected negative program responses for each exploratory endothelial priority: interferon-gamma response, inflammatory response and complement for ETS2; and IL-6/JAK-STAT3 signaling, interferon-gamma response and TNF-alpha/NF-kappaB signaling for BHLHE40. Points are donor-equal mean scores, intervals are donor-bootstrap 95% intervals, and exact sign-test P and within-knockout q values are reported separately from global GSEA FDR. Figure 6f retained genes that belonged to the same TF-program GSEA leading edge in both tissues, had negative predicted effects in both, and passed gene-level Benjamini-Hochberg FDR < 0.05 in both tissue models. Direct fitted-network targets and propagated responses were distinguished in source data; neither category establishes experimental binding or causal regulation.

**SUPPLEMENTARY FIGURES S9 AND S10 ORGANIZATION**

Supplementary Figure S9a reports network support and knockout estimability for all 18 endothelial candidates. Burgundy symbols indicate muscle support and blue-gray symbols brain support; filled symbols indicate sufficient support, hollow symbols indicate one to four fitted edges, and gray marks indicate no fitted edges or an unavailable perturbation. Supplementary Figure S9b displays donor-equal source-program responses, and S9c reports the cross-tissue evidence audit. Supplementary Figure S9d is the complete 18-candidate vector atlas: 15 candidates have maps in at least one tissue, whereas PLSCR1, RBCK1 and TRAFD1 are retained as gray non-estimable placeholders. Placement within S9d has no prescribed order and does not indicate rank.

Supplementary Figure S10a provides the muscle and early-CPS-window brain myeloid reference state maps and contains no knockout result. Supplementary Figure S10b reports tissue-specific network support using the same burgundy muscle and blue-gray brain convention, S10c displays donor-equal source-program responses, and S10d reports the program-level robustness audit. Supplementary Figure S10e is one combined atlas containing virtual-knockout and randomized-residual maps for IRF1, STAT3, SPI1 and STAT2, with gray non-estimable placeholders for MXD1 and PLSCR1. Placement within S10e has no prescribed order and does not indicate rank.

**INTERPRETATION BOUNDARIES**

CellOracle results remain conditional on the observational expression data, promoter prior, selected features and fitted networks. Cross-tissue agreement supports exploratory candidate prioritization and does not establish pathway inhibition in vivo, endothelial or myeloid state rescue, direct muscle-to-brain signaling, or therapeutic efficacy. CPS windows are cross-sectional analytical ranges rather than longitudinal disease stages, and donors or participants, rather than cells, nuclei or display grid points, are the biological replicates unless explicitly stated otherwise.

**Supplementary Table S1. Characteristics of the analyzed muscle and brain donors**

| **Dataset / analysis subset** | **n** | **Age range (years)** | **Median age (years)** | **Female / male, n** |
| --- | --- | --- | --- | --- |
| HLMA Young | 8 | 15–45 | 31.5 | 3 / 5 |
| HLMA Old | 15 | 77–99 | 82 | 8 / 7 |
| SEA-AD full atlas | 84 | 65–102 | 90 | 51 / 33 |
| SEA-AD inference subset | 73 | 69–102 | 91 | 43 / 30 |
| SEA-AD early CPS window | 30 | 72–102 | 90 | 16 / 14 |
| SEA-AD late CPS window | 48 | 69–100 | 91 | 30 / 18 |
| SEA-AD severely affected (excluded from inference) | 11 | 65–96 | 90 | 8 / 3 |

Source metadata were summarized for the analyzed released objects. Ages are recorded ages; donor counts, not nuclei, define sample size. HLMA Young and Old are age-defined groups, not clinical sarcopenia diagnoses. The 73-donor SEA-AD inference subset excludes the 11 source-designated severely affected donors. The early (CPS ≤ 0.55) and late (CPS > 0.45) windows overlap in five donors and must not be summed as independent groups. Recorded no-dementia/dementia counts were 41/32 in the inference subset, 25/5 in the early window, 19/29 in the late window and 1/10 among excluded donors. AD neuropathological change counts (not AD / low / intermediate / high) were 9/12/21/31, 9/11/8/2, 0/4/15/29 and 0/0/0/11, respectively. CPS is a cross-sectional neuropathology-derived ordering and is not equivalent to MCI. PBMC and bulk-blood cohort sizes and their distinct statistical units are reported in Methods.
