## Supplemental_File_2_Supplementary_Figures for "Cross-Tissue Inflammatory Convergence Supports a Putative Muscle–Brain Axis in Alzheimer’s Disease"

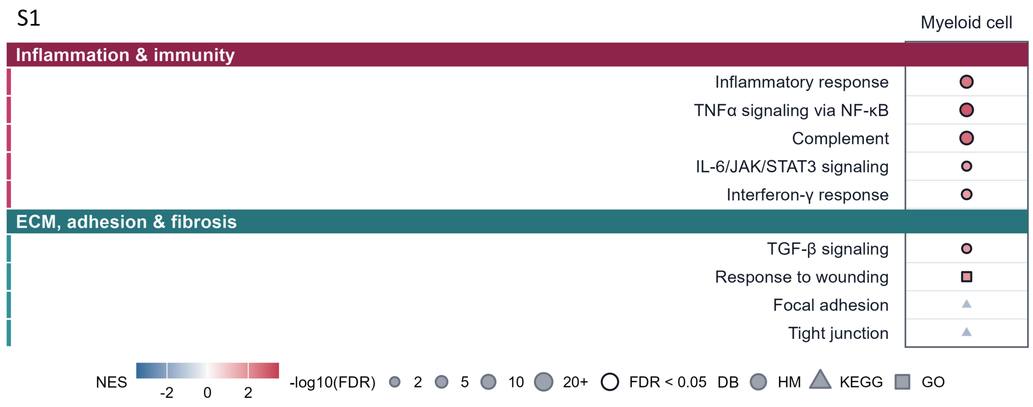


**Supplementary Fig. S1 Myeloid MAST-GSEA pathways aligned with the Figure 1 modules.** Nine of the 16 pathways displayed in Fig. 1e met MAST-GSEA nominal P < 0.05 in muscle myeloid cells: five inflammatory or immune and four extracellular-matrix, adhesion, or fibrosis pathways. The cohort comprised 4 Young donors (259 nuclei) and 9 Old donors (4,166 nuclei). Positive NESs favor Old. Dot size represents −log10(FDR), and dark outlines denote original database-wide Benjamini–Hochberg FDR < 0.05. Hallmark, KEGG Legacy, and Gene Ontology Biological Process collections retain their original correction families. Focal adhesion and tight junction were nominally significant but did not meet FDR < 0.05. Nominal P values reaching 10−8 reflect the computational floor. Donors, rather than individual nuclei, are the biological replicates.

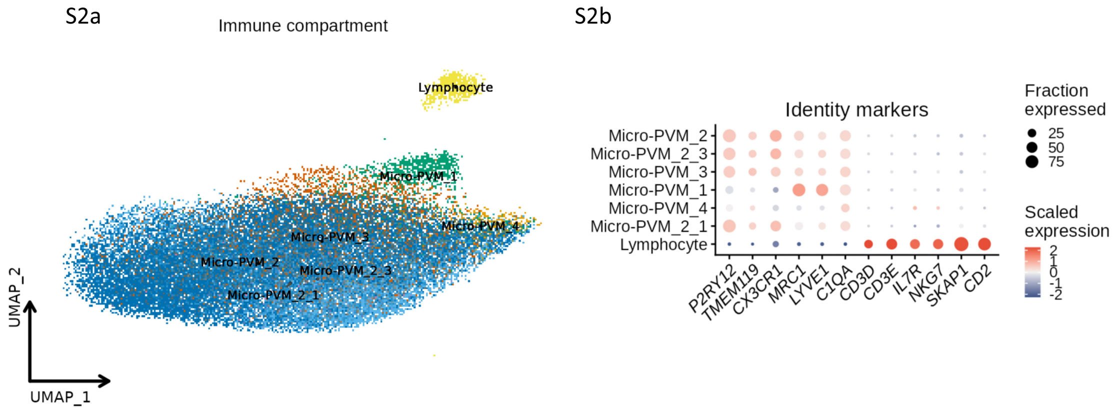


**Supplementary Fig. S2 SEA-AD immune-cell identities and analytical scope.** Left, local view of the original SEA-AD middle temporal gyrus immune UMAP, retaining 36,674 nuclei after exclusion of 12 Monocyte nuclei. Six Microglia-PVM supertypes and Lymphocyte are shown in the original embedding. Right, expression of 12 identity markers; dot size denotes the expressing fraction and color the gene-scaled mean expression. Downstream Micro-PVM analyses pooled the six myeloid supertypes and excluded Lymphocyte and Monocyte. This figure documents cell identity and analytical scope and does not present a between-group test.

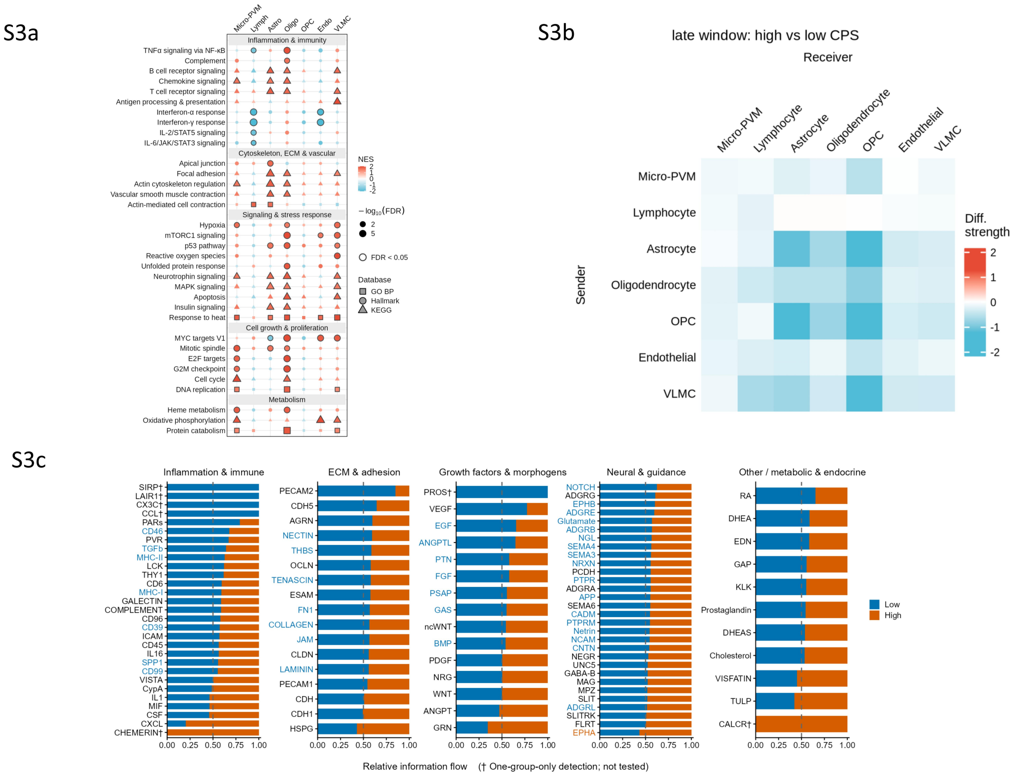


**Supplementary Fig. S3 Late-window transcriptional programs and inferred communication in SEA-AD.** Top left, GSEA of 34 pathways across seven non-neuronal populations in the late CPS window (CPS > 0.45; 48 donors). Positive NESs denote enrichment with increasing CPS; dark outlines indicate original FDR < 0.05. The Lymphocyte model included 45 donors; other models included 48. Top right, CellChat High-minus-Low communication differences after a within-window median split (24 donors per group). Bottom, relative information flow for 102 pathways; Low and High shares sum to 1 within each pathway. Two-sided paired Wilcoxon tests compared matched ligand–receptor records, with Benjamini–Hochberg adjustment across 95 tested pathways. Daggers identify seven pathways dtected in one group only and therefore not tested. Five donors overlapped the early and late windows; these are cross-sectional analytical ranges rather than independent groups or longitudinal stages. Communication analyses are pooled model inferences, not donor-level tests.

S4

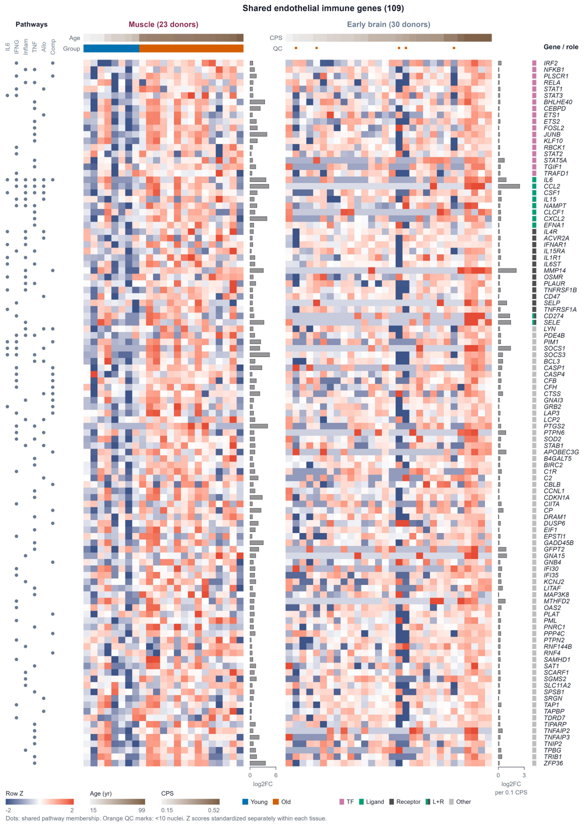


**Supplementary Fig. S4 Complete shared endothelial leading-edge genes across muscle and brain.** All 109 genes in the union of shared leading edges from the six Fig. 3 pathways are shown, including the 41 annotated genes in the main panel and 68 other genes. Muscle comprised 23 donors (8 Young, 15 Old; 23,227 endothelial cells), and brain comprised 30 donors (693 endothelial nuclei). Columns are ordered by age or CPS. Gene-wise Z scores were calculated separately within each tissue and were not used for cross-tissue clustering. Dot tracks denote shared pathway membership; role annotations indicate TF, ligand, receptor, dual identity, or other. Five brain donors with fewer than 10 endothelial nuclei are flagged. Side bars show muscle limma Old-versus-Young log2 fold changes and brain NEBULA log2 fold changes per 0.1 CPS. Display of the complete set adds no gene-level significance filter; “other” does not imply absence of biological function.

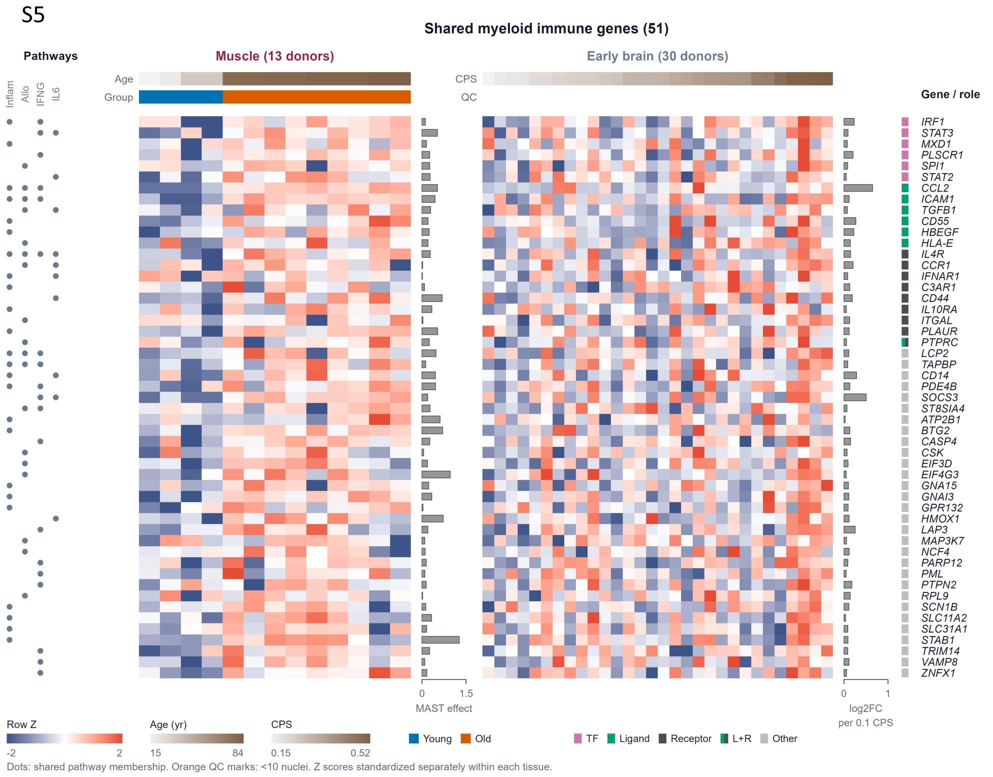


**Supplementary Fig. S5 Complete shared myeloid leading-edge genes across muscle and brain.** All 51 genes in the intersection of the four Fig. 4 pathways are shown, including 21 annotated genes and 30 other genes. Muscle comprised 4 Young donors (259 nuclei) and 9 Old donors (4,166 nuclei); brain comprised 30 donors (13,416 Micro-PVM nuclei). Columns are ordered by age or CPS, and gene-wise Z scores were calculated separately within each tissue. Dot tracks denote shared pathway membership, and role annotations follow Supplementary Fig. S4. Muscle side bars show MAST-adjusted predicted mean log2-expression differences; brain bars show NEBULA log2 fold changes per 0.1 CPS. The complete display does not imply that every gene was differentially expressed at FDR < 0.05 in both tissues and does not constitute independent-cohort validation.

S6

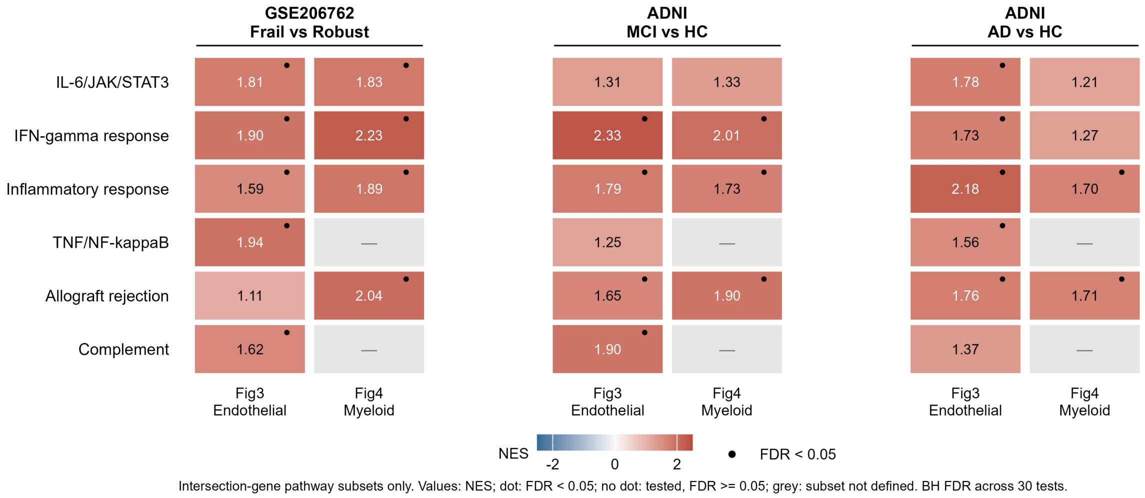


**Supplementary Fig. S6 Bulk enrichment of endothelial and myeloid intersection-gene pathways.** Fixed gene subsets contained only the Fig. 3 endothelial or Fig. 4 myeloid muscle–brain intersection genes assigned to six source-related Hallmark pathways. Values are NESs from preranked GSEA of the complete eligible bulk-gene ranking. GSE206762 compares 10 Frail and 10 Robust RNA pools; ADNI compares MCI or AD with HC in a joint 732-participant model. Dots denote FDR < 0.05. Unmarked colored tiles were tested but did not meet this threshold. Gray tiles indicate that no Fig. 4 myeloid intersection-gene subset was defined for complement or TNF-α/NF-κB signaling and do not denote nonsignificant tests. Benjamini–Hochberg correction was applied across 30 prespecified source-subset–pathway tests. Positive NESs indicate enrichment toward higher expression in the case group.

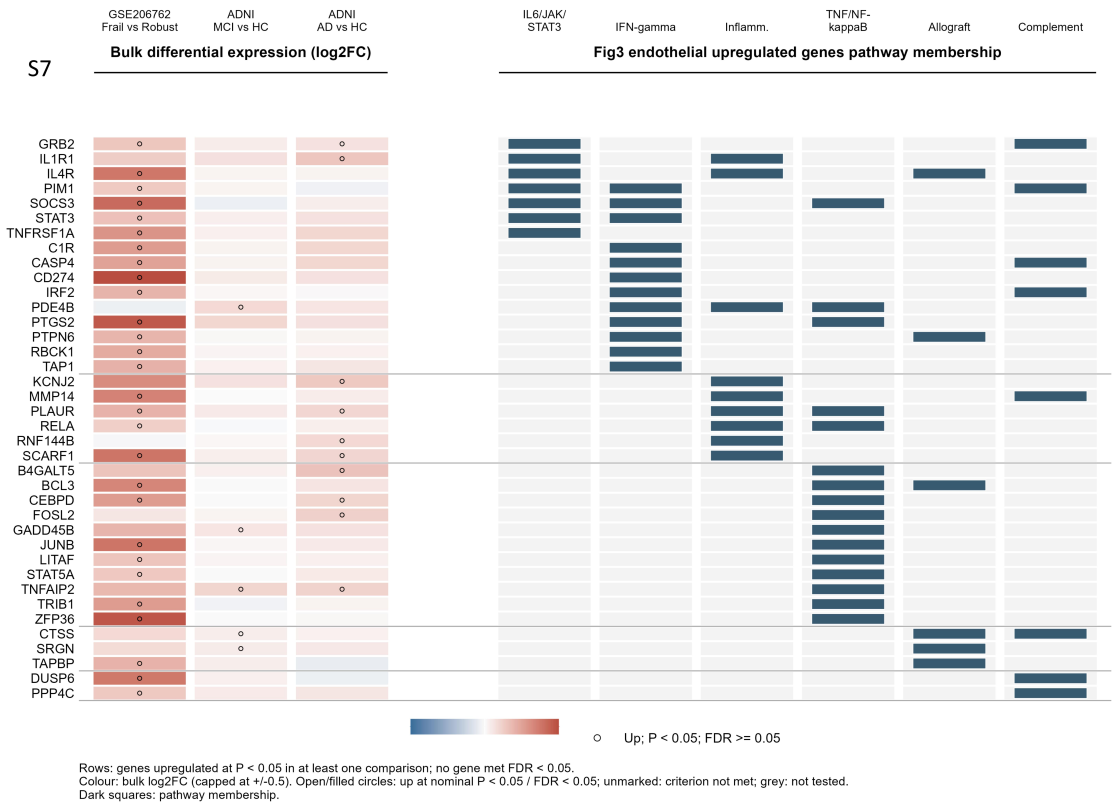


**Supplementary Fig. S7 Bulk projection of upregulated endothelial intersection genes.** Rows show the 38 of 109 Fig. 3 endothelial genes with log2 fold change > 0 and nominal P < 0.05 in at least one of GSE206762 Frail versus Robust, ADNI MCI versus HC, or ADNI AD versus HC. The three left columns show effect estimates for every available comparison, including nonsignificant or opposite-direction estimates for selected genes. Open circles denote comparison-specific upregulation at P < 0.05 and FDR ⩾ 0.05; no displayed gene met gene-level FDR < 0.05. Gray indicates an unavailable test. Dark squares indicate fixed membership in six source pathways; genes may belong to multiple pathways. The display scale is capped at ±0.5, whereas exact effects, P values, and FDR values are retained in the source table.

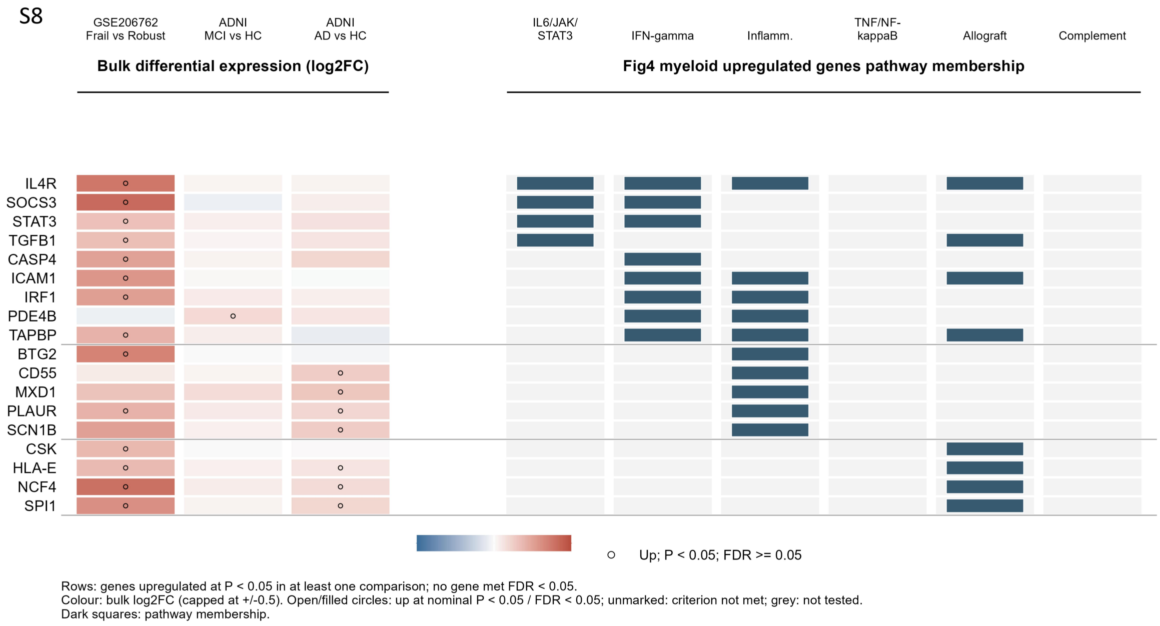


**Supplementary Fig. S8 Bulk projection of upregulated myeloid intersection genes.** Rows show the 18 of 51 Fig. 4 myeloid genes with log2 fold change > 0 and nominal P < 0.05 in at least one bulk comparison. Encoding, comparisons, and significance definitions follow Supplementary Fig. S7. No displayed gene met gene-level FDR < 0.05. Fig. 4 provided no intersection genes assigned to complement or TNF-α/NF-κB signaling; the corresponding membership columns therefore contain no marked genes.

S9a

**Network support for virtual KO**

Filled: KO support (burgundy muscle; blue-gray brain); M/B boxes: tissue estimable

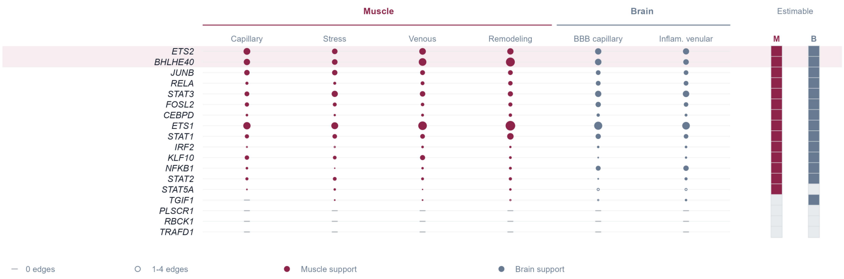


S9b

**Predicted source-program responses**

Color: donor-equal scaled response; dot: conditional 95% CI below zero

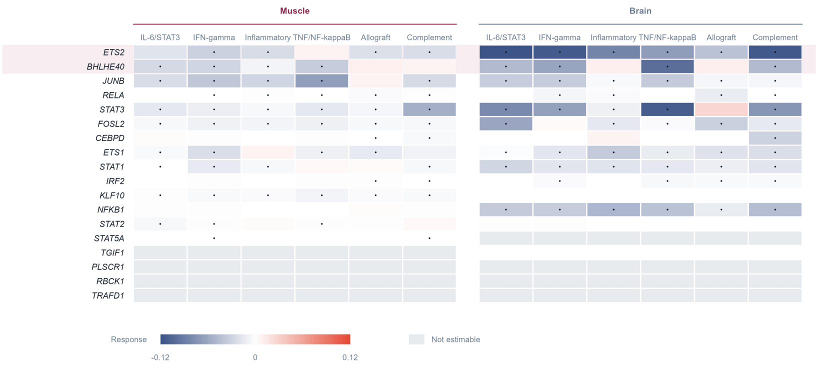


90

S9c

### Cross-tissue evidence ranking

Thirteen TFs estimable in both tissues; cells count qualifying source programs, out of six

| | Negative both | 95% CI <0 both | 5/5 refits both | 6/6 variants both | snRNA/tech consistent | NES <0 both | Stable negative | Evidence priority | GSEA q<0.05 both | Full rule |
|---|---|---|---|---|---|---|---|---|---|---|
| **ETS2** | 5 | 4 | 4 | 5 | 6 | 5 | 3 | 3 | 0 | 0 |
| **BHLHE40** | 3 | 3 | 3 | 4 | 6 | 4 | 3 | 3 | 0 | 0 |
| *JUNB* | 5 | 5 | 3 | 5 | 5 | 5 | 3 | 2 | 0 | 0 |
| *RELA* | 5 | 4 | 2 | 2 | 4 | 5 | 2 | 2 | 0 | 0 |
| *STAT3* | 5 | 5 | 1 | 3 | 5 | 5 | 1 | 1 | 0 | 0 |
| *FOSL2* | 5 | 5 | 2 | 5 | 5 | 4 | 1 | 1 | 0 | 0 |
| *CEBPD* | 1 | 1 | 1 | 2 | 2 | 3 | 1 | 1 | 0 | 0 |
| *ETS1* | 5 | 4 | 0 | 3 | 5 | 2 | 0 | 0 | 0 | 0 |
| *STAT1* | 4 | 4 | 3 | 2 | 4 | 4 | 0 | 0 | 0 | 0 |
| *IRF2* | 2 | 2 | 0 | 0 | 5 | 3 | 0 | 0 | 0 | 0 |
| *KLF10* | 0 | 0 | 0 | 0 | 1 | 5 | 0 | 0 | 0 | 0 |
| *NFKB1* | 0 | 0 | 2 | 1 | 1 | 1 | 0 | 0 | 0 | 0 |
| *STAT2* | 0 | 0 | 0 | 0 | 4 | 0 | 0 | 0 | 0 | 0 |

ETS2 and BHLHE40 are tied at three evidence-priority programs; no program passes the full rule.

91

S9d

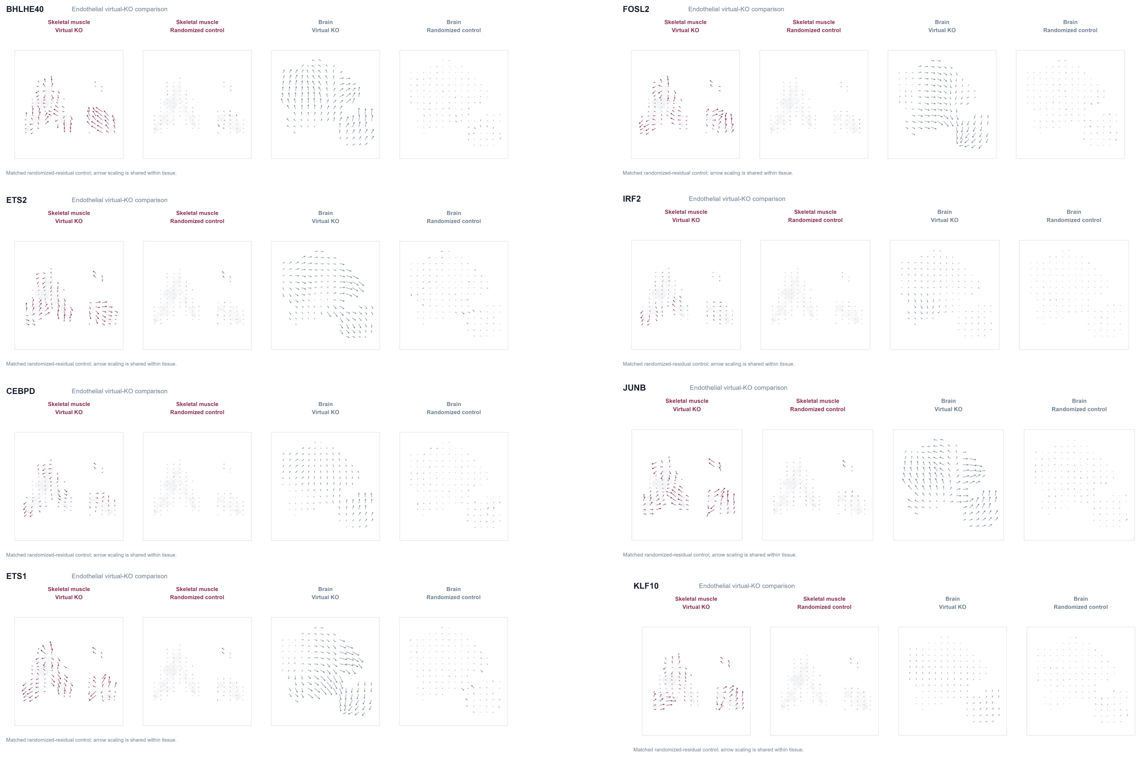

BHLHE40
Endothelial virtual-KO comparison
Skeletal muscle Virtual KO
Skeletal muscle Randomized control
Brain Virtual KO
Brain Randomized control
Matched randomized-residual control; arrow scaling is shared within tissue.
FOSL2
ETS2
IRF2
CEBPD
JUNB
ETS1
KLF10


92

S9d

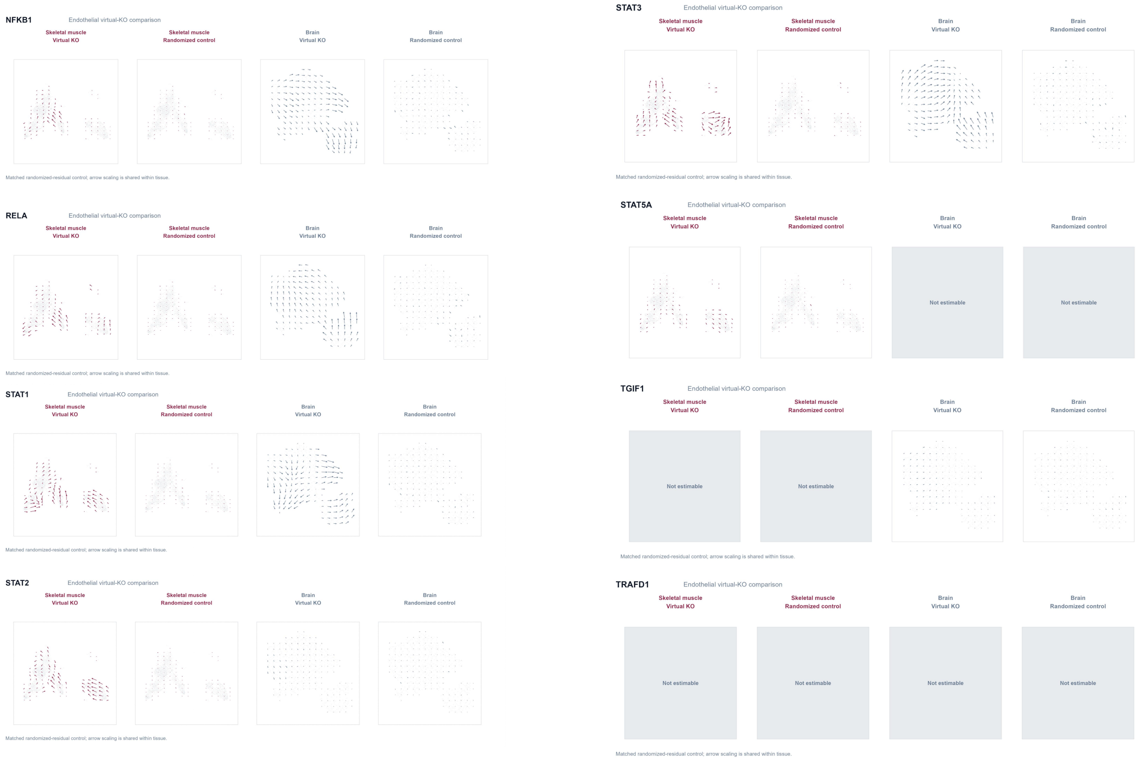

NFKB1
Endothelial virtual-KO comparison
Skeletal muscle Virtual KO
Skeletal muscle Randomized control
Brain Virtual KO
Brain Randomized control
Matched randomized-residual control; arrow scaling is shared within tissue.
STAT3
RELA
STAT5A
Not estimable
STAT1
TGIF1
STAT2
TRAFD1


93

S9d

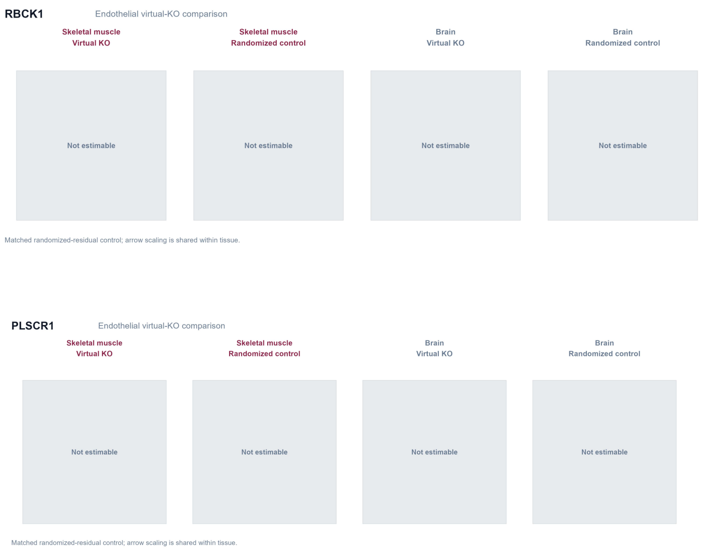


**Supplementary Fig. S9 Cross-tissue virtual-knockout screen of endothelial transcription factors. (a)** Network support and whole-tissue estimability for virtual KO of all 18 endothelial TF candidates. Dot area scales with fitted outgoing TF–target edges. Burgundy filled circles denote muscle states meeting the KO support rule; blue-gray filled circles denote supported brain states. Hollow circles indicate only 1–4 fitted edges and insufficient KO support; gray dashes indicate no fitted edges. M and B boxes denote whole-tissue estimability. This panel reports model feasibility rather than KO effect, direction, or significance. **(b)** Donor-equal standardized responses of six source inflammatory programs after KO in muscle and brain endothelial cells. Color denotes response direction and magnitude; dots mark conditional 95% intervals below zero; gray denotes non-estimable combinations. **(c)** Cross-tissue evidence audit for 13 TFs estimable in both tissues. Cells report the number of six programs meeting each robustness criterion. *ETS2* and *BHLHE40* each had three evidence-priority programs, the highest count, but no program met the complete global-FDR rule. **(d)** Virtual-KO and randomized-residual control maps for the 15 candidates estimable in at least one tissue:

*ETS2, BHLHE40, JUNB, RELA, STAT3, FOSL2, CEBPD, ETS1, STAT1, IRF2, KLF10, NFKB1, STAT2, STAT5A,* and *TGIF1*. TF placement within d has no prescribed order and does not indicate rank. *PLSCR1, RBCK1,* and *TRAFD1* were not estimable in either tissue and are retained as gray placeholders in the individual files. Arrow scaling is shared within tissue but not across tissues. Minimum global GSEA q values were 0.989 in muscle and 0.978 in brain; ranking reflects directional robustness, not confirmed pathway suppression.

S10a

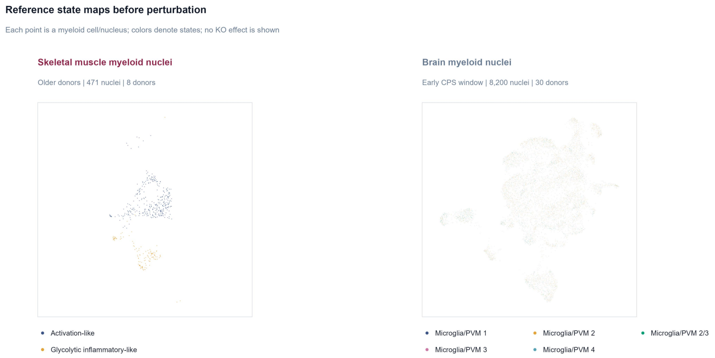


S10b

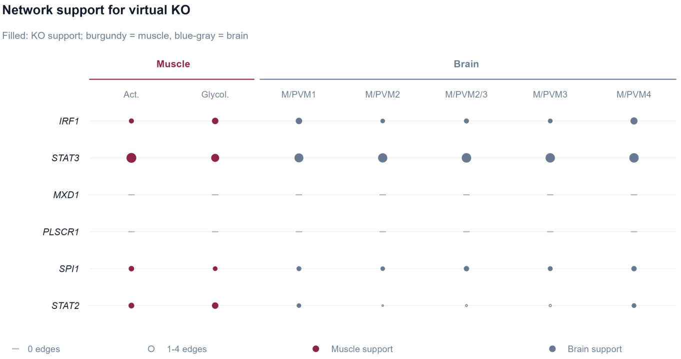

S10c

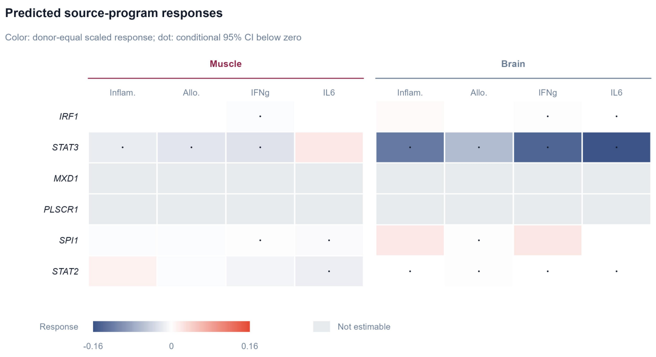


S10d

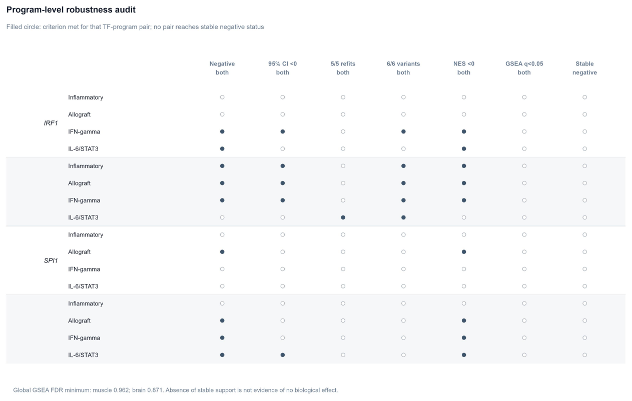


S10e

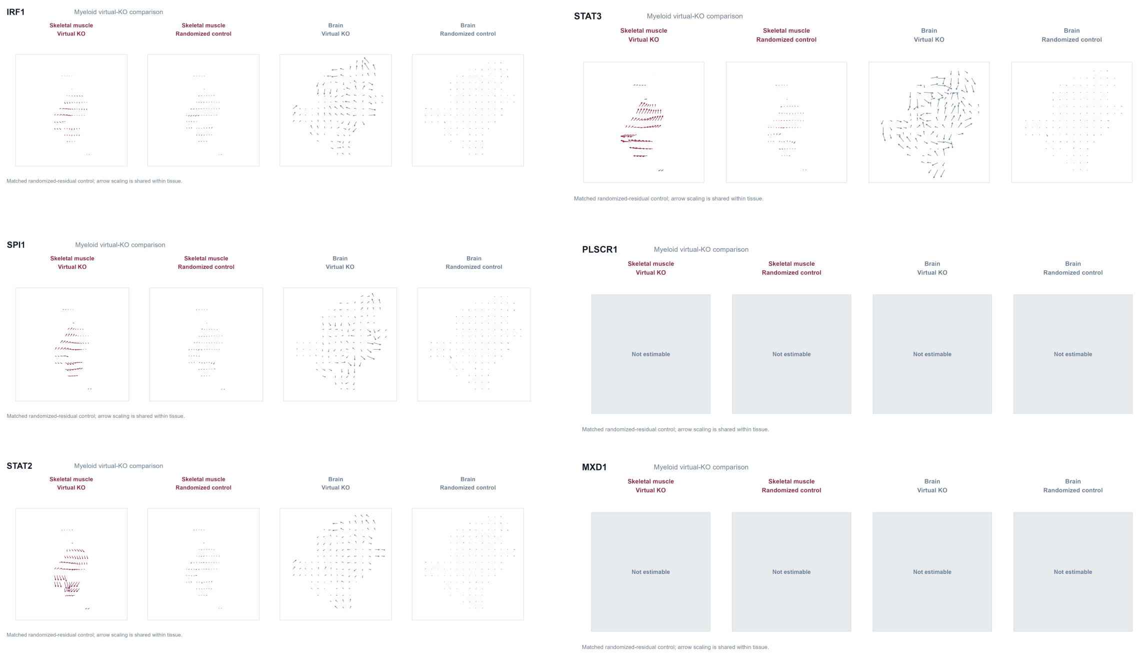

**Supplementary Fig. S10 Virtual-knockout analysis of myeloid transcription factors.** **(a)** Reference myeloid state spaces before perturbation. Muscle comprised 471 single-nucleus profiles from eight older donors and two marker-informed provisional states; brain comprised 8,200 nuclei from 30 early-CPS-window donors and five retained Microglia-PVM source states. This panel provides state context and contains no KO result. **(b)** Network support for virtual KO of six myeloid TF candidates. Burgundy filled circles denote supported muscle states and blue-gray filled circles supported brain states; hollow circles indicate 1–4 fitted edges and gray dashes no fitted edges. IRF1, STAT3, SPI1, and STAT2 were estimable; MXD1 and PLSCR1 were not. This panel reports feasibility rather than perturbation magnitude, direction, or significance. **(c)** Donor-equal standardized responses of four Fig. 4 source programs after KO in muscle and brain myeloid cells; encoding follows Supplementary Fig. S9b. **(d)** Program-level robustness audit for the four estimable TFs. No TF–program pair reached stable-negative status or global GSEA $q < 0.05$ in both tissues; minimum global q values were 0.962 in muscle and 0.871 in brain. Failure to meet this rule does not imply absence of biological function. **(e)** Combined virtual-KO and randomized-residual control atlas for IRF1, STAT3, SPI1, and STAT2, with gray non-estimable placeholders for MXD1 and PLSCR1. TF placement within e has no prescribed order and does not indicate rank. Scaling is shared within tissue but not across tissues. CellOracle results are predictions from observational single-cell data and do not establish causal rescue.
